# Aging, dauer, and stature phenotypes are conferred by structure-directed missense mutations in the endogenous AGE-1/phosphatidylinositol 3-kinase catalytic subunit

**DOI:** 10.64898/2026.01.15.699774

**Authors:** You Wu, Tam Duong, Neal R. Rasmussen, Kent L. Rossman, David J. Reiner

**Affiliations:** College of Medicine, Texas A&M Health Science Center, Texas A&M University, Houston, TX 77030, USA; Institute of Biosciences and Technology, Texas A&M Health Science Center, Texas A&M University, Houston, TX 77030, USA; Division of Abdominal Transplantation, School of Medicine, University of North Carolina, Chapel Hill, NC 27599, USA

**Keywords:** phosphatidylinositol 3-kinase, AGE-1, LET-60/Ras, lifespan, aging, animal size, dauer diapause, vulval precursor cells (VPCs)

## Abstract

Phosphatidylinositol 3-kinase (PI3K) integrates insulin/IGF signaling (IIS) and Ras inputs to control lifespan, metabolism and growth, yet the organismal consequences of selective structural perturbations remain poorly understood. Using structure-guided CRISPR/Cas9-dependent genome editing, we dissected functions of AGE-1, the sole Class IA PI3K catalytic subunit in *Caenorhabditis elegans*. An endogenously tagged AGE-1, containing a long flexible linker, epitope and fluorescent tag, retained full activity, enabling visualization of native protein dynamics *in vivo*. A constitutively active E630K substitution, modeled on oncogenic p110α alleles, markedly shortened lifespan and enhanced Ras-dependent induction of primary vulval precursor cell (VPC) fate, confirming evolutionary conservation of PI3K activation mechanisms that directly modulate longevity and development. Structural modeling further guided mutation of AGE-1 residues predicted to mediate Ras binding. Unexpectedly, a putative AGE-1 variant defective in Ras association, together with a complementary Ras effector-binding mutation, produced enlarged animals with reduced dauer formation. These phenotypes reveal a previously unrecognized Ras>PI3K signaling axis that restrains somatic growth and promotes entry into diapause, counter to canonical IIS models. Together, these structure-informed alleles show that discrete PI3K structural perturbations can differentially uncouple lifespan, growth, and developmental outcomes *in vivo*. By combining structural modeling with genome editing in a tractable aging model, this work establishes a framework for dissecting conserved signaling enzymes at single-residue resolution and uncovers unexpected organismal roles for PI3K structure in coordinating growth and longevity.

## Introduction

Research across diverse systems – including human cohorts, rodent models, cell-based assays, *in vitro* biochemistry/biophysics/reconstitution, computer modeling, and invertebrate organisms – can yield insights greater than the sum of their parts. Each platform has distinct strengths and limitations, and findings are often most powerful when integrated across disciplines. This perspective highlights the importance of what might be called “model discipline”: knowing where a model excels, where it falls short, and how technological advances may shift those boundaries.

Mammalian cell-based systems permit rapid manipulations such as RNA interference (RNAi), or CRISPR/Cas9-mediated depletion, often in high-throughput formats. However, they are often limited by incomplete knockdown of endogenous proteins and gene redundancy, often with 2-4 paralogs per group in mammals (Dehal & Boore, 2005). Mouse models can provide physiological relevance for human disease and lifespan but are costly and slow, complicating mechanistic linkage between protein structure and phenotype. Biochemical and biophysical studies precisely define molecular interactions but are typically divorced from organismal physiology.

Invertebrate models can bridge these gaps. *Caenorhabditis elegans* combines genetic tractability with the complex biological organization of a multicellular animal. “The Worm” is transparent and highly amenable to transgenesis and genome editing. With 959 somatic cells, 302 neurons, and a fully mapped cell lineage and neural connectome, *C. elegans* enables precise, *in vivo* analysis of conserved signaling networks and their impact on biology. These physical attributes are complemented by relatively little paralog redundancy while retaining broad conservation with disease-relevant genes (Kim, Underwood, Greenwald, & Shaye, 2018). The short generation time (∼2.5 days), maximum lifespan (∼18 days), and ease of culturing make *C. elegans* an ideal animal for exploration of mechanistic underpinnings of aging and lifespan while including a sophisticated genetic toolkit. Not surprisingly, discoveries in *C. elegans* have repeatedly advanced understanding of metabolism and aging (Kropp, Bauer, Zafra, Graham, & Golden, 2021; Murphy & Hu, 2013), beginning with the identification of *age-1*, the first genetic determinant of lifespan in any system (Johnson, 1990; Klass, 1983).

Biochemical insights often drive genetic experiments in *C. elegans*. For example, catalytic residues defined in mammalian kinases and exchange factors for small GTPases can be tested *in vivo* to probe biological effects and functional conservation of proteins (Shin et al., 2019; Shin, Kaplan, Duong, Fakieh, & Reiner, 2018). When sequence conservation is limited, structural information becomes particularly valuable: three-dimensional folds, domain interfaces, and higher-order protein relationships are often preserved even when primary sequence identity diverges. Structural modeling thus enables rational design of targeted mutations for genome editing in model organisms.

We apply this approach to the catalytic subunit of Class IA phosphatidylinositol 3-kinase (PI3K), encoded in mammals by PIK3CA, PIK3CB, and PIK3CD to produce p110α, p110β, and p110δ (hereafter PI3Kcat). PI3Kcat converts phosphatidylinositol (4,5)-bisphosphate (PtdIns(4,5)P_2_, or PIP_2_) to phosphatidylinositol (3,4,5)-trisphosphate (PtdIns(3,4,5)P_3_, or PIP_3_), which in turn activates downstream kinases PDK1 and AKT/PKB (Engelman, Luo, & Cantley, 2006; Fruman et al., 2017). In *C. elegans*, the sole PI3Kcat ortholog, AGE-1, acts centrally within the conserved insulin/IGF-like signaling (IIS) cascade, together with DAF-2/InsR, AAP-1 (regulatory PI3K subunit), IST-1 (IRS1 adaptor), DAF-18/PTEN, PDK-1, and AKT-1/2, to regulate lifespan, dauer diapause, and metabolic adaptation, processes central to human health (Murphy & Hu, 2013; Partridge, Alic, Bjedov, & Piper, 2011). Notably, these components were identified by classical forward genetic screens, rather than via structure-guided hypotheses (Hu, 2007). But the flourishing of CRISPR as an editing tool positions us to dissect functions of the AGE-1 complex with a scalpel rather than a hammer.

PI3Kcat is activated by two inputs: direct recruitment to phosphotyrosines on the insulin/insulin-like growth factor (IGF) receptor (InsR), a receptor tyrosine kinase (RTK), and by binding of GTP-bound Ras (Rodriguez-Viciana et al., 1994). Both mechanisms are conserved, and their balance influences downstream signaling and physiological outcomes. Structural studies in *Drosophila* and mouse PI3Kcat revealed residues required for Ras binding (Gupta et al., 2007; Orme, Alrubaie, Bradley, Walker, & Leevers, 2006), but their functional significance at the organismal level has not been tested in a system where lifespan and metabolic state can be directly assayed.

Here, we use structure-guided CRISPR/Cas9-dependent genome editing to dissect PI3Kcat function *in vivo*. We engineered into endogenous AGE-1 protein (1) a C-terminal fluorescent protein+epitope tag with a long, flexible linker to enable live imaging without impairing function; (2) a gain-of-function substitution modeled on constitutively active p110α alleles from cancer, to test effects on lifespan and insulin/EGF growth factor-related signaling; and (3) a mutation in the Ras-binding domain (RBD), guided by structural modeling of Ras-PI3Kcat interface. Unexpectedly, the RBD mutation increased body size and decreased dauer formation – phenotypes recapitulated by a complementary effector-binding mutation in the effector-binding loop of LET-60/Ras – revealing a Ras>PI3K axis that restrains somatic growth, contrary to canonical IIS predictions.

Together, these results demonstrate how structural modeling can inform genome editing to produce precise, interpretable alleles that probe conserved signaling mechanisms *in vivo*. More broadly, this approach bridges molecular structure and organismal physiology, providing a framework for understanding how defined PI3K perturbations influence aging, metabolism, and developmental plasticity.

## RESULTS

### Endogenous AGE-1/PI3Kcat is expressed ubiquitously with no evident subcellular localization

The domain organizations of AGE-1/PI3Kcat, AAP-1/PI3KR, and LET-60/Ras are shown (**Fig. 1A**). To visualize the expression of AGE-1 in live animals, we used CRISPR/Cas9-dependent genome editing to insert sequences encoding mNeonGreen fluorescent protein (mNG) and 2xHA epitope tag into the endogenous *age-1* gene. First, threading of the *C. elegans* sequence of AGE-1/PI3Kcat, particularly the positioning of the C-term ⍺-helix 5 (⍺5) of LET-60, the N-term of AGE-1, and the positioning of the PIP2 head group (**Fig. 1B**; **S1A,B**), suggest that an N-terminal tag may interfere with the association of AGE-1 with the plasma membrane. Second, we observed an N-terminal extension in *C. elegans* AGE-1: 57 residues longer than human PIK3CA and 38 residues longer than *Drosophila* PI3K92E. This AGE-1 extension contains seven Arginine residues relative to one in *Drosophila* (**Fig. S1C**). Based on location in the protein relative to the plasma membrane and established mechanisms of proteins associating with the plasma membrane, this extension may constitute an electrostatic interaction between the N-terminus of AGE-1 and acidic microdomains in the plasma membrane that is not present in orthologous proteins, thus counter-indicating an N-terminal tag in them. Third, C-terminal tagging may sterically hinder AGE-1 protein function. Publication of structures of human PI3Kα and KRAS during preparation of this manuscript corroborated these observations (Czyzyk et al., 2025; Torosyan et al., 2025).

**Figure 1.**
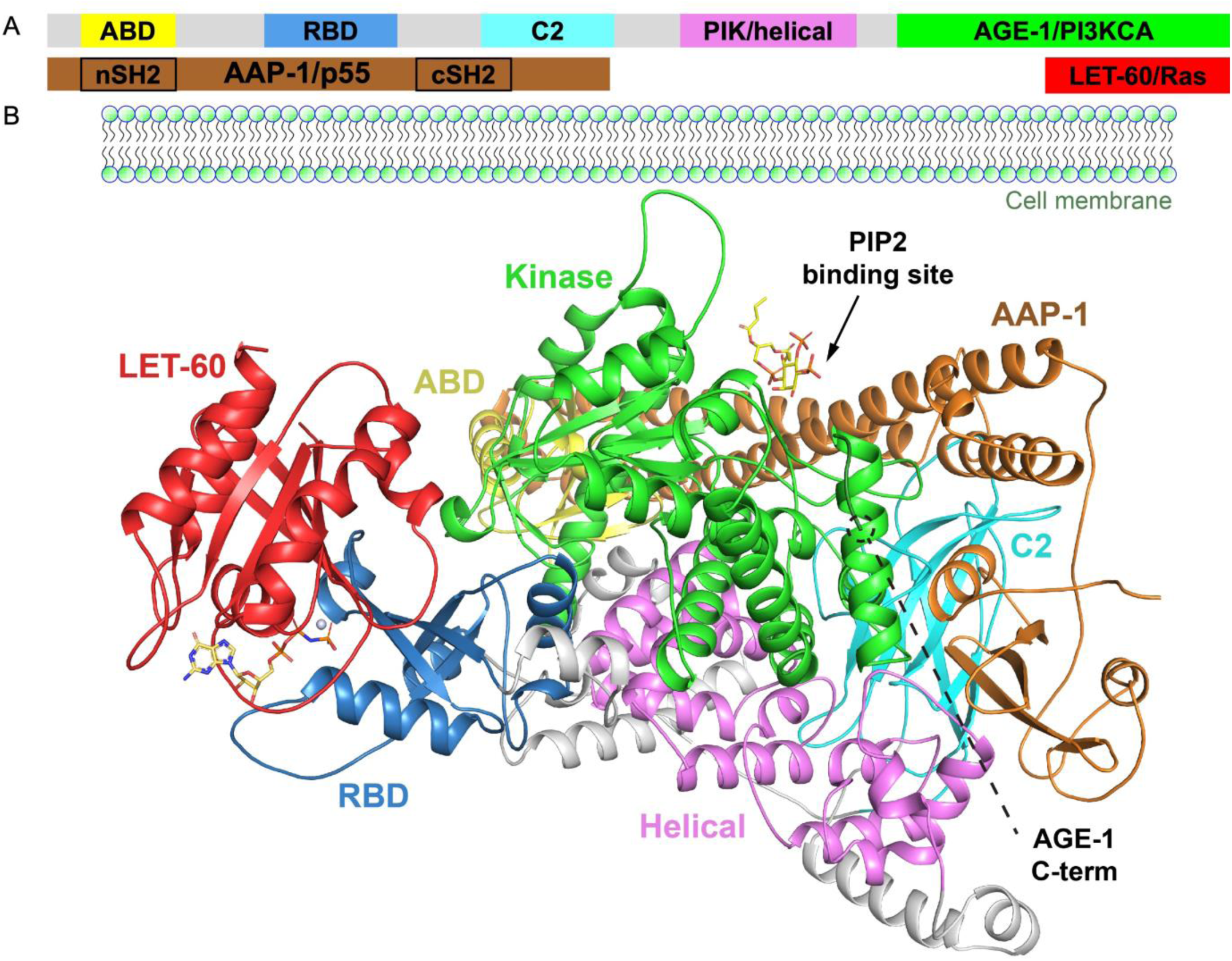
A structural model of the Ras - PI 3-Kinase signaling complex. **(A)** Conserved domains diagram of AGE-1/PI3Kcat, AAP-1/p55/PI3Kreg and LET-60/Ras small GTPase. Domains from left to right: ABD = <u>a</u>daptor <u>b</u>inding <u>d</u>omain; RBD = <u>R</u>as <u>b</u>inding <u>d</u>omain; C2 = PI(4,5)P_2_ binding; PIK/helical = conserved in PI3Ks and PI4Ks; PI3KCA = catalytic lipid kinase domain; SH2 = phosphotyrosine binding domain associated with binding p-Tyr in signaling cascades. AGE-1: **<u>age</u>**ing alteration. Catalytic subunit of PI3K (PI3KCA). AAP-1: <u>A</u>d<u>ap</u>tor for PI3K. p55/p85 in mammals. Note that *C. elegans* AAP-1 does not encode an identifiable inter-SH2 sequence (iSH2) domain found in mammalian p50/p55 regulatory subunits, but a coiled-coil is detected in the central region, which like substitutes for the same activity and the structural level. LET-60, **<u>let</u>**hal; The Ras small GTPase orthologous to human H,N,K-Ras. (**B)** A *post hoc* structural model of the predicted *C. elegans* AGE-1 PI3KCA catalytic subunit, a portion of the AAP-1 ortholog of the human p50/p55 regulatory subunit (not including cSH2, which is also absent for various mammalian structures) and LET-60 with Mg^++^ and GMPPNP, a non-hydrolysable GTP analog. The plasma membrane is oriented upwards in the diagram; accordingly, the C-terminal end of LET-60 ⍺5 is oriented upwards. (The C-terminal hypervariable region+CAAX is truncated in most structures since the sequence is unstructured and lipid modified. But the HVR is expected to abut the PM and the CAAX (Cys-Ali-Ali-X, where Ali = aliphatic and X = any residue) is modified with a farnesyl lipid and proteolytically processed.) The putative PI(4,5)P_2_ binding sequence and phosphoinositol headgroup, shown, is also upwardly oriented toward the expected location of the PM. See Methods for assembly of this structural model. This C-term is indicated, located far enough from the PM to potentially avoid steric conflict with the association of the enzyme complex with the PM. The CRISPR/Cas9-dependent insertion was at the C-term, with a 30-residue linker consisting of 10xGAS, followed by mNeonGreen (mNG) and 2xHA epitope tag.

Consequently, to minimize the likelihood of perturbing AGE-1 function, we inserted sequences encoding a 30-residue linker at the 3’ end of *age-1* but 5’ to the sequences encoding mNG::2xHA (**Fig. 2A**). The resulting *age-1(re353[age-1::linker::mNG::2xHA])* mutant, here referred to as the *age-1(re353[tag])* mutant animal, was detected by triplex PCR and validated by sequencing (see Methods and see **Table S2** for oligonucleotide sequences.)

**Figure 2.**
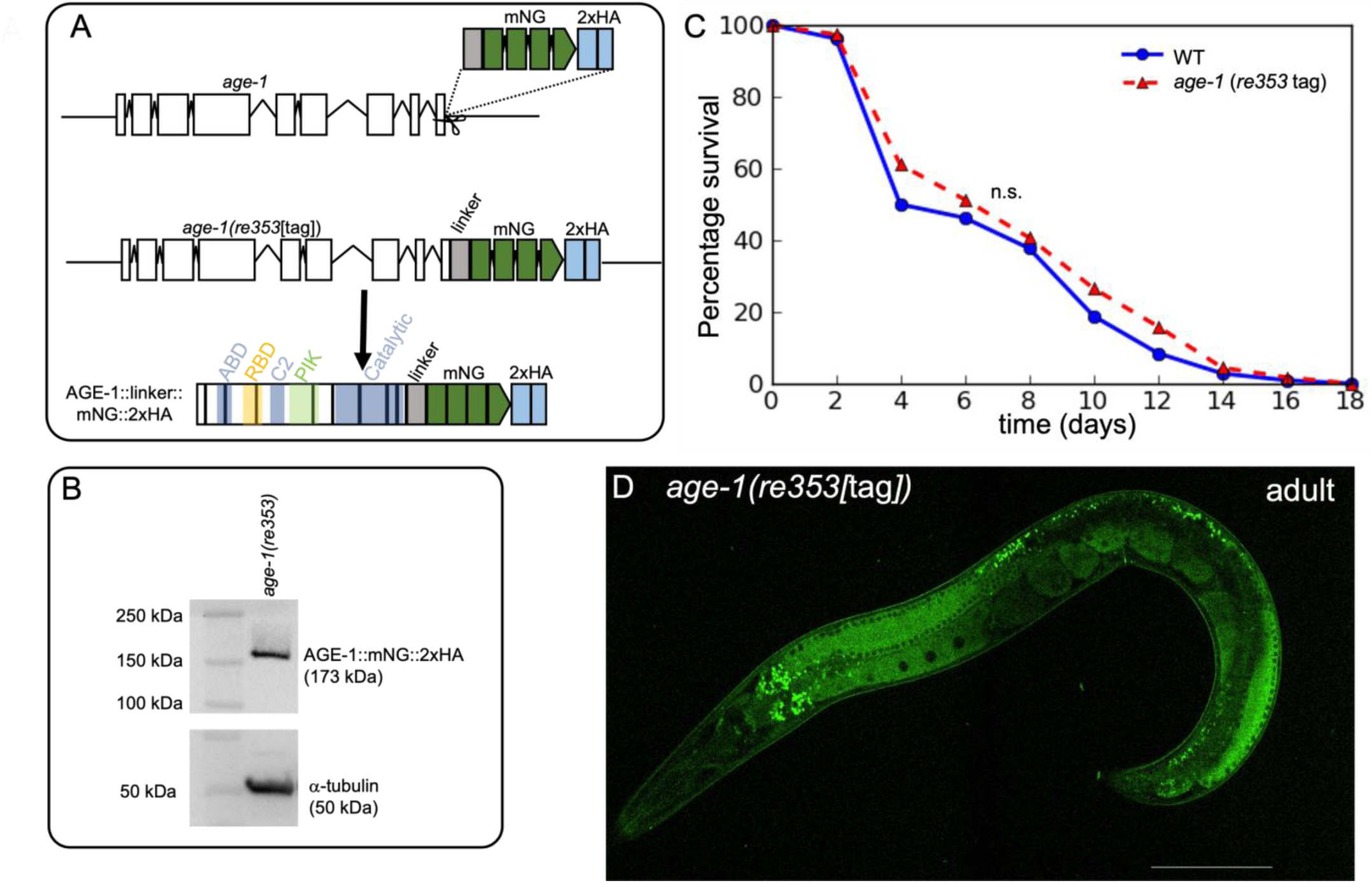
Tagging endogenous AGE-1 at the C-terminus. **(A)** A schematic of the *age-1* gene and 3’-end insertion strategy for a C-terminally tagged endogenous protein. The exon-intron gene, defined by extensive RNAseq data, is shown at the top with the PCR product repair template used, generated with 35 bp homology arms at each end to support homology-directed repair (HDR). The resulting construct (middle) was detected and tracked by triplex PCR and confirmed by Sanger sequencing of all novel and flanking sequences. The bottom model shown the predicted resulting protein; colors indicate different functional domains. Lines indicate different exons. **(B)** Immunoblotting of edited *age-1(re*353[tag]) animals with ⍺-FLAG antibody revealed the predicted 173 kDA protein predicted by AGE-1+tag sequences, with ⍺-tubulin control. **(C)** Kaplan-Meier curves show percent survival of wild-type animals *vs. age-1(re353*[tag]) animals, analyzed concurrently, with no difference observed. N = sample size. P-value was calculated by Log-rank Mantel-Cox test. n.s. = no significant difference. **(D)** A spinning disk confocal photomicrograph (488 nm) of an optical section of an *age-1(re*353[tag]) adult animal, showing likely ubiquitous expression. Left = anterior, down = dorsal. Dark circles are nuclei, particularly standing out in oocytes in the proximal gonad and small nuclei in the distal gonad. Bright punctae are intestinal autofluorescence of gut granules, which are lipid storage depots. Additional images are shown in (**Fig. S2**). Variability of intensity depends on position in the low-magnification field with variable excitation. Scale bar is 100 μm.

After western blotting, immunodetection using anti-HA antibodies revealed a band in *age-1(re353[tag])* animals consistent with the 173 kDa expected from AGE-1::linker::mNG::2xHA (**Fig. 2B**). To assay impact of tagging on the function of AGE-1, we assayed lifespan of *age-1(re353[tag])* animals vs. the wild type. The two lifespans were not significantly different (**Fig. 2C**).

We surveyed expression of AGE-1::linker::mNG::2xHA in *age-1(re353[tag])* animals using spinning disk confocal microscopy. Expression was ubiquitous in the animal with potentially elevated signal in the germline (**Fig. 2D**). Expression was cytoplasmic, as evinced by nuclear exclusion of AGE-1. Expression was also consistent throughout developmental stages, including embryos (**Fig. S2A-D**). We did not observe evidence of subcellular localization, though we only examined this phenomenon closely in the vulval precursor cells (**Fig. S2C**), when a role for AGE-1 signaling had been hypothesized (see below). Thus, we observed that AGE-1/PI3Kcat is expressed globally, with each cell throughout the life of the animal potentially capable to respond to upstream signals to activate AGE-1 activity.

### Constitutively activated endogenous AGE-1/PI3Kcat decreases lifespan consistent with established roles of other IIS components

From a very large cohort, 12.1% of tumors harbor activating missense mutations in human PI3Kcat/p110α, encoded by PI3KCA (Sivakumar et al., 2023; Zhao & Vogt, 2008). These cluster in a hotspot in the helical/PIK domain N-terminal to the lipid kinase domain (**Fig. 3B**; **S3A,B**). These mutations alter either of the first two Glutamates (Es), E542 or E545, of the four acidic residues in the “**E**IT**E**QEKD” sequence, changing them to Lysines (K) in a charge reversal. Oncogenic mutations in this region are thought to disrupt the salt-bridge inhibitory interaction with the SH2 domain of the p55/p85 regulatory subunit of PI3K (Carson et al., 2008; Zhao & Vogt, 2008). Alignment with *Drosophila* and *C. elegans* AGE-1/PI3Kcat orthologs with human PI3Kα reveals that the helical/PIK domain is not well conserved at level of amino acid sequence, yet we observe three and four acidic residues in this interval, respectively, in human PI3Kcat/p110α (**Fig. 3B**; **S3A,B**). In *C. elegans* this sequence is clustered as “VL**EE**DEQ”. Threading *C. elegans* sequences onto the mammalian structure reveal similar structural positioning of *C. elegans* E630 and human E545 (**Fig. 3A**). Thus *C. elegans* AGE-1 could plausibly be activated by an E630K mutation.

**Figure 3.**
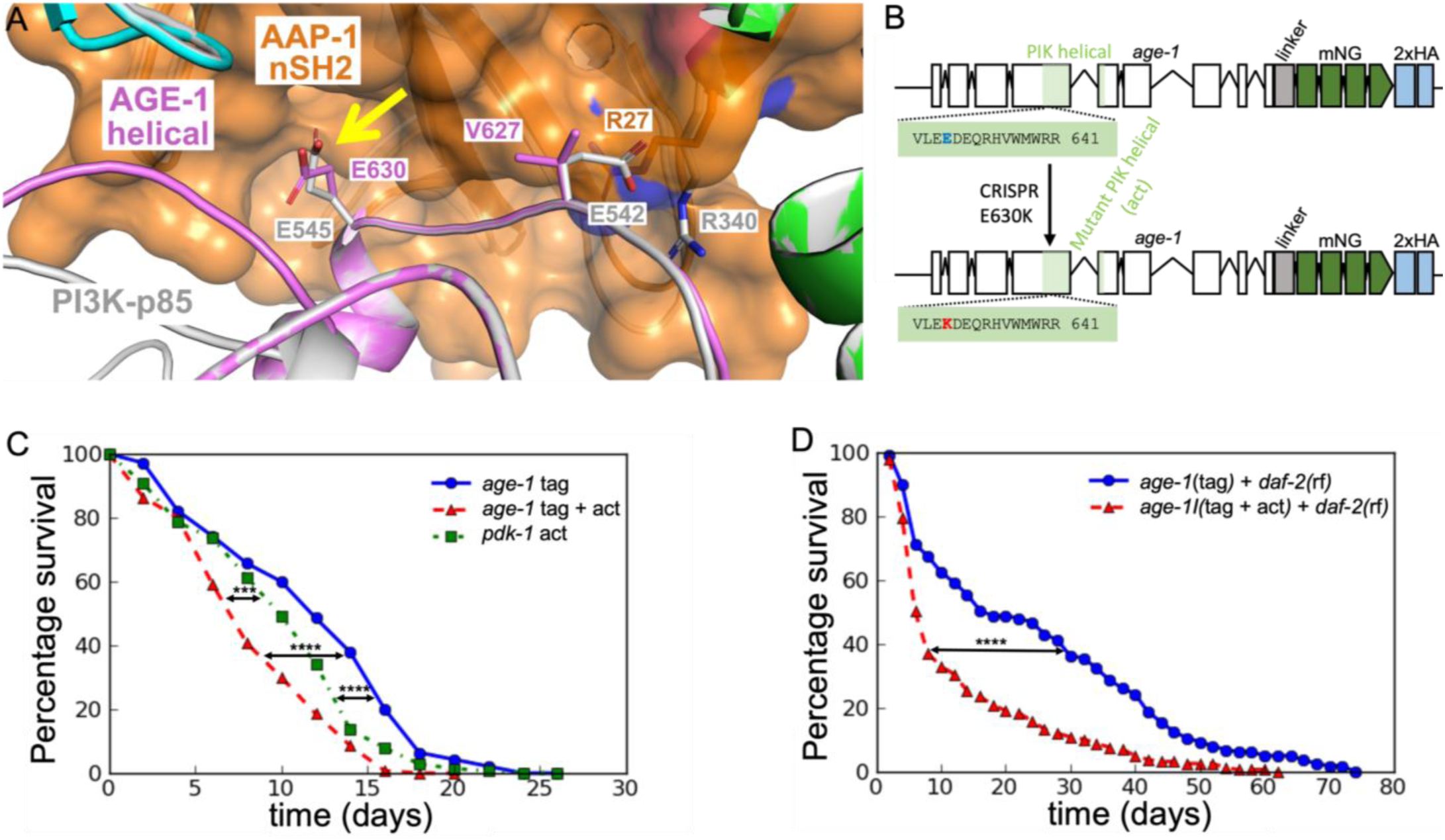
The *C. elegans* E630K equivalent to human oncogenic mutation E545K causes a gain of function. **(A)** A zoom of a structural model of the inhibitory interaction of the helical/PIK domain of PI3K catalytic subunit with the nSH2 domain of the regulatory subunit (pink and brown, as in Fig. 1). Threaded AGE-1-AAP-1 structure is overlaid with the PI3Kcat-p85 structure, human-nematode. The yellow arrow indicates the overlay depicts of human E545 (gray) and nematode E630 (pink). Solid pink indicates predicted AGE-1, solid gray depicts solid human PI3K, mottled pink/gray indicates predicted co-positioning of the two polypeptides. While the E545/E630 residues align, the human E542, also commonly mutated to a K in human cancers, aligns with nematode V627 (see sequence alignments in **Fig. S3A,B**). **(B)** A schematic of *age-1(re353[age-1::linker::mNG::2xHA])* with the wild-type E630 residue above in blue and the putative gain-of-function E630K mutation below in red, encoded by *age-1(re353re392*gf[tag E630K]). **(C)** A Kaplan-Meier curve showing that *age-1(re353re392*gf[tag E630K]) and *pdk-1(mg142*gf) gain-of-function mutants for the PI3K>PDK>Akt pathway live shorter than *age-1(re353[tag])* animals, as predicted by previous genetic analysis. Arrows indicate P values. N = 140 animals for each genotype. **(D)** Activating mutant *age-1(re353re392*gf[tag E630K]) also strongly reverses the longevity conferred by reduced function of InsR *daf-2(e1370*rf) with *age-1(re353[age-1::linker::mNG::2xHA*]), thus with only the E630K missense mutation between the strains. N= 140 for *age-1(re353*[tag]*); daf-2(e1370rf)*, N = 160 for *age-1(re353re392gf*[tag E630K]*); daf-2(e1370rf)*. P values calculated by Kaplan-Meier estimator, Log-rank (Mantel-Cox) test. For p values, * ≤ 0.05, ** ≤ 0.01, *** ≤ 0.001, **** ≤ 0.0001.

To introduce a potentially constitutively activating mutation into our endogenous AGE-1::linker::mNG::2xHA, we used CRISPR/Cas9-dependent genome editing to alter E630 to K, thereby generating *age-1(re353re392*gf*[tag E630K])*. Immunoblotting with anti-HA antibody revealed no change in stability in the tag E630K AGE-1 protein relative to *age-1(re353[tag])* alone (**Fig. S3C**).

The *age-1(hx546)* reduction-of-function mutation in *C. elegans* PI3Kcat was the first lifespan-extending mutation found in animals (Friedman & Johnson, 1988a, 1988b; Johnson, 1990; Klass, 1983). We reasoned that a constitutively activating, gain-of-function mutation in AGE-1 would reduce lifespan, consistent with gain-of-function mutations in PDK-1 and AKT-1 (Paradis, Ailion, Toker, Thomas, & Ruvkun, 1999; Paradis & Ruvkun, 1998) and loss of DAF-18/PTEN lipid phosphatase function (Ogg & Ruvkun, 1998). We measured the lifespan of *age-1(re353re392*gf*[tag E630K])* animals compared to that of *age-1(re353[tag])* animals and the published gain-of-function *pdk-1(mg142*gf) mutant animals (Paradis et al., 1999). As expected, *pdk-1(mg142*gf) (“*pdk-1* act”) animals lived shorter than *age-1(re353[tag])* animals, which lived a wild-type span (**Fig. 3C**). *age-1(re353re392*gf*[tag E630K]) (*“*age-1* tag + act”) animals lived shorter than not only *age-1(re353[tag])* animals but also *pdk-1(mg142*gf) animals (**Fig. 3C**). The *age-1(re353re392*gf*[tag E630K])* mutation also reversed the longevity conferred by mutation of DAF-2/InsR, *daf-2(e1370)* (**Fig. 3D**). This epistatic interaction is consistent with the established biochemical role of PI3K downstream of Insulin/IGF receptors in signal transduction (Dorman, Albinder, Shroyer, & Kenyon, 1995). These results indicate that *age-1(re353re392*gf*[tag E630K])* confers a strong lifespan defect and is likely constitutively active signaling protein.

### Constitutively activated endogenous AGE-1/PI3Kcat increases induction of 1°vulval precursor cell fate

In response to a point source of epithelial growth factor (EGF) from the ventral gonad, the initially equipotent *C. elegans* vulval precursor cells (VPCs) are induced to form the 3°-3°-2°-1°-2°-3° pattern of cell fates with 99.8% accuracy (Braendle & Felix, 2008; Shin et al., 2019; **Fig. 4A**). The necessary and sufficient signaling cascades that direct this developmental event are the EGFR>Ras>Raf>MEK>ERK canonical MAP kinase pathway promoting 1° cell fate and the DSL>Notch receptor>CSL pathway promoting 2° cell fate (Shin et al., 2018). Two modulatory pathways also contribute to this process: PDK-1/PDK>AKT-1/Akt promotes 1° fate in support of the canonical ERK/MAP kinase pathway while the EGFR>Ras>RalGEF>Ral cascade promotes 2° cell fate in support of the Notch pathway (Shin et al., 2019; Shin et al., 2018). (3° fate is promoted by an independent Rap2>MAP4K signal (Fakieh & Reiner, 2025).) An inhibitory role for DAF-18/PTEN has been described for the 1°-promoting modulatory signal (Nakdimon, Walser, Frohli, & Hajnal, 2012; Shin et al., 2019); gain of AGE-1/PI3Kcat function would be expected to phenocopy loss of DAF-18/PTEN

**Figure 4.**
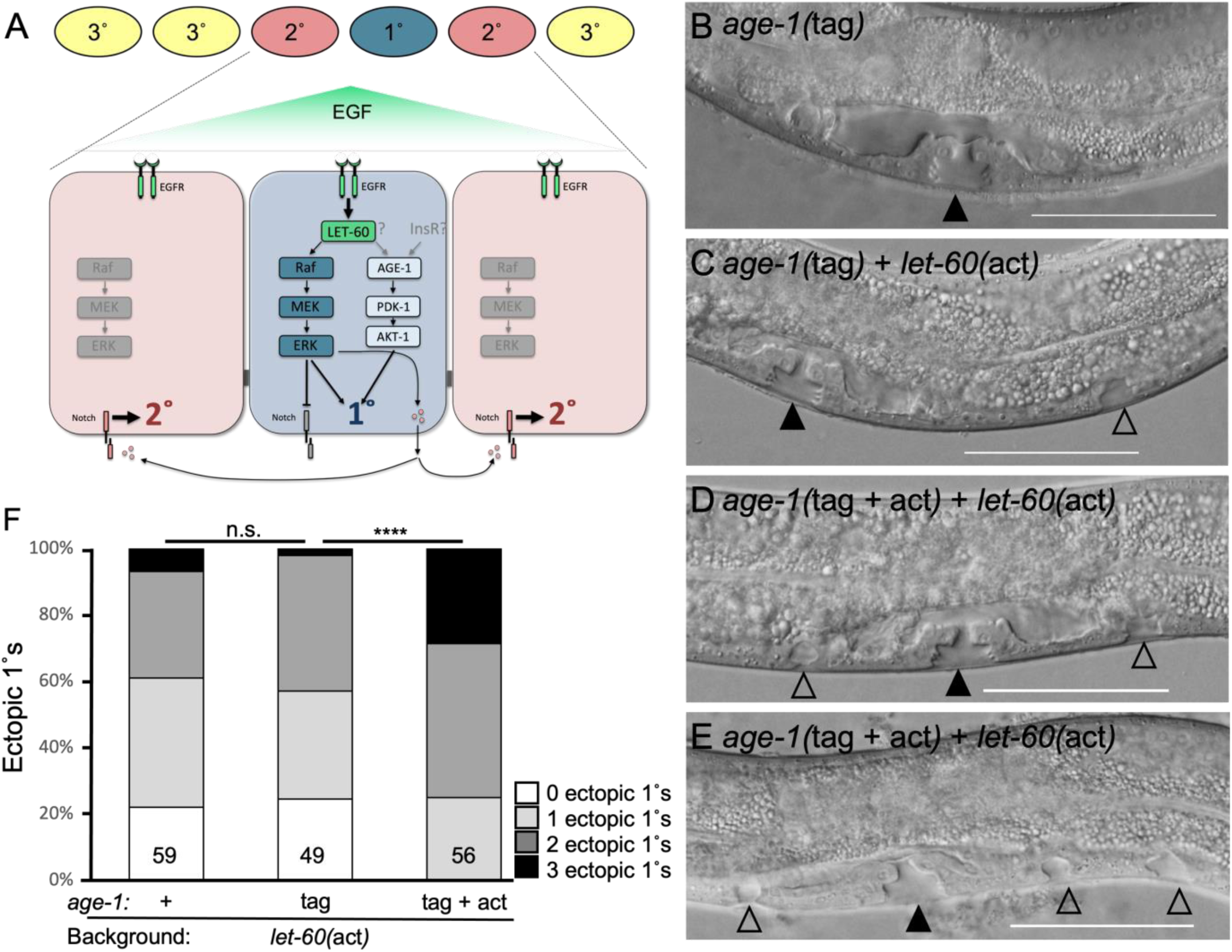
Constitutive activation of AGE-1/PI3Kcat increases induction of 1° cell fates among VPCs. **(A)** A schematic of developmental patterning of the vulval precursor cells (VPCs). EGF signaling induces 1° fate via the necessary and sufficient EGFR>Ras>Raf>MEK>ERK/MAP kinase pathway. PDK->AKT-1 promotes 1° fate as a modulatory pathway, with inhibition by DAF-18/PTEN lipid phosphatase implying a potential 1°-promoting role of AGE-1/PI3Kcat. The source of activating signal, DAF-2/InsR, or LET-60/Ras, is unknown. **(B-E)** Photomicrographs of DIC imaging of late 4^th^ larval stage (L4) animals with zero **(B)** to three **(E)** ectopic 1° lineages. Black arrowheads indicate the normal morphogenetic vulva, formed of 2°-1°-2° lineages. Open arrowheads indicate ectopic pseudovulvae, formed from inappropriately induced 1° lineages. Genotypes are in black. *let-60(n1046*gf) is a G13E mutant that moderately induces ectopic 1° cells and is sensitive to increases or decreases to 1°-promoting signaling. Ventral is down and anterior is left. Scale bars = 50 µm. **(F)** Quantification of ectopic 1° pseudovulvae from each genotype. N is indicated in the bar for each genotype. P value was calculated by the Mann-Whitney test.

The VPC patterning system is significantly buffered against extraneous developmental noise (Braendle & Felix, 2008; Shin et al., 2019). Consequently, the impacts of genetic perturbations of these modulatory signals are most easily detected using the modestly constitutively activating G13E mutation in the *C. elegans* Ras ortholog, *let-60(n1046*gf) (Fakieh & Reiner, 2025; Rasmussen, Dickinson, & Reiner, 2018; Shin et al., 2019; Shin et al., 2018; Zand, Reiner, & Der, 2011). In this strain, increases or decreases in induction of ectopic 1° pseudovulvae indicate decreases or increases in 1°-promoting activity, respectively (**Fig. 4A-E**). AGE-1::linker::mNG::2xHA, encoded by *age-1(re353[tag])*, did not alter the induction of ectopic 1° pseudovulvae by *let-60(n1046*gf) compared to no mutation of *age-1* (**Fig. 4F**; **S4A**). This observation is consistent with the observation that tagged AGE-1 also does not alter function, as measure with lifespan. In contrast, *age-1(re353re392*gf*[tagE630K])* strongly increased induction of ectopic 1° pseudovulvae (**Fig. 4F**; **S4A**), thus phenocopying loss of DAF-18/PTEN and gain of PDK-1 and AKT-1 functions.

Taken together, our results with *age-1(re353re392*gf*[tagE630K])* reducing lifespan and promoting formation of ectopic 1° pseudovulvae indicate that E630K constitutively activates endogenous AGE-1/PI3Kcat. The efficacy of this mutation validates our structure-based approach for tagging endogenous invertebrate proteins and introducing mutations likely to phenocopy pathogenic mutations found in mammalian systems.

### Reduction-of-function mutations in the Ras>AGE-1/PI3Kcat binding interface reveal a role in restricting animal stature

Activity of the PI3K catalytic subunit can be activated through two general mechanisms: i) recruitment of PI3Kcat to the receptor tyrosine kinase at the plasma membrane via the p55/p85 regulatory subunit, or ii) direct activation via binding of GTP-bound Ras to the Ras binding domain (RBD) of PI3Kcat and recruitment to the plasma membrane (Cuesta, Arevalo-Alameda, & Castellano, 2021). A single mutation, *age-1(ag12),* introducing an L336F change in the RBD of AGE-1 (L267F in human PI3KCA), was isolated previously in a mutant screen for increased resistance to infection by the pathogen *Pseudomonas aeruginosa* (Miyata, Begun, Troemel, & Ausubel, 2008). This L336F change of *age-1(ag12)* also increased animal lifespan and at 25°C caused formation of dauer rather than L3 larvae, a diapause arrested alternative third larval stage whose induction is frequently associated with low IIS activity. Moreover, this L336F mutant residue lies in the hydrophobic interior of the RBD domain; the increased size of the L336F side chain may thus disrupt hydrophobic packing of the interior and potentially destabilize the RBD or the entire AGE-1 protein. Consequently, the L336F mutation may not specifically disrupt binding of LET-60/Ras to the RBD of AGE-1. (A mutant strain containing the *age-1(ag12)* could not be recovered from the original strain collection.)

Consequently, we undertook a structure-directed approach to disrupt the binding interface of LET-60/Ras and the RBD of AGE-1. The core effector binding loop of LET-60/Ras is 100% conserved among model invertebrates and mammals, presumably from the evolutionary constraints of a single short polypeptide required to interact with several effector proteins (Reiner & Lundquist, 2018). In contrast, the primary sequence of the AGE-1 RBD is not well conserved with the RBD of mammalian PI3K catalytic subunits (**Fig. S5A,B**). Consequently, we used structural models to infer the importance of two basic residues, Arg-303 and Lys-304, predicted to regulate and lie the binding interface, respectively, of LET-60/Ras and the AGE-1/PI3Kcat RBD (**Fig. 5A**).

**Figure 5.**
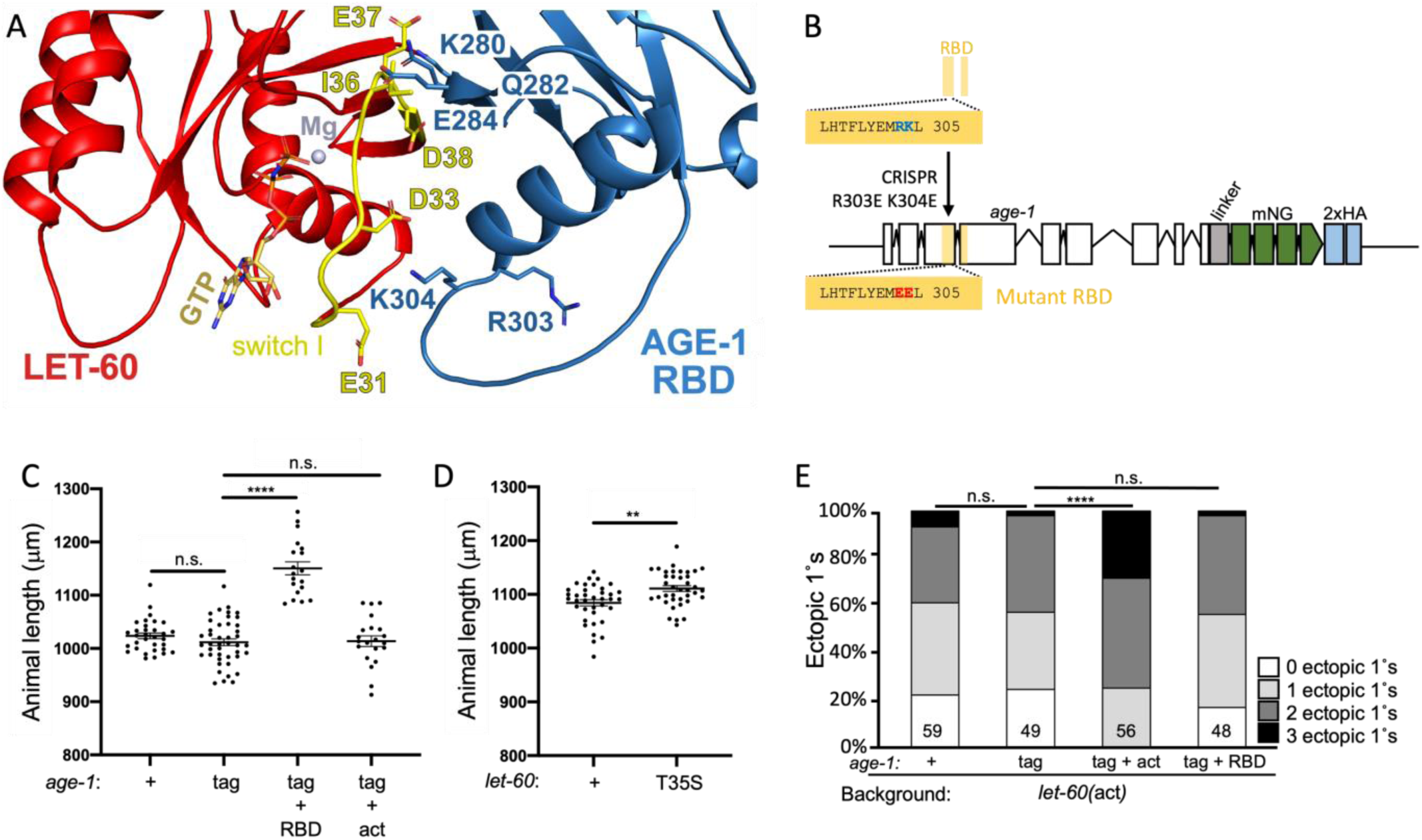
Defective association of LET-60/Ras with AGE-1/PI3Kcat reveals a role in constraining body stature. **(A)** A zoomed portion of the same structural model as in Fig. 1B of LET-60/Ras and the RBD of AGE-1/PI3Kcat. K304 of the AGE-1 RBD may interact with Switch I of LET-60/Ras, such that replacement with the charge-reversing side chain with Glu (E) may repel LET-60 and reduce binding. R303 faces the neighboring loop, such that replacement with the charge-reversing side chain with Glu (E) may interfere with positioning of the interface of the RBD. Also shown are K280, Q282, and E284 which are also predicted to disrupt the interface with LET-60 if mutated to E, A, and K, respectively, in some combination. But the sgRNA for CRISPR for R303 and K304 was predicted to be more robust, so we mutated that site first and were successful. **(B)** A schematic of *age-1(re353[age-1::linker::mNG::2xHA])* with the wild-type R303 K304 residues above in blue and the R303E K304E putative interface disrupting mutation below in red, encoded by *age-1(re353re377*rf[tag R303E K304E]). **(C)** Animal length in microns as measured from DIC photomicrographs for RBD and activating mutations in AGE-1. (**D)** Animal length in microns as measured from DIC photomicrographs for the LET-60 T35S putative Raf-selective mutant vs. wild type. **(E)** Quantification of ectopic 1° pseudovulvae in the *let-60(n1046*gf) background reveals no role of the RBD in 1° fate induction. n.s. = not significant, ** < 0.01, **** 0.0001, P value was calculated by the *t*-test (**C**,**D**) and Mann Whitney test (**E**).

We used CRISPR/Cas9-dependent genome editing to substitute codons encoding acidic Glutamate residues for the wild-type basic Arg-303 and Lys-304 codons in *age-1(re353[tag])*, resulting in the *age-1(re353re377*rf*[age-1::mNG::2xHA(tag R303E,K304E)])* mutant animal. The R303E K304E mutation did not destabilize AGE-1::linker::mNG::2xHA protein as detected by immunoblotting for HA (**Fig. S5C**).

Unexpectedly, we observed that animals were of greater length than stage-matched animals encoding wild-type or tagged AGE-1. This was true for both animal length (**Fig. 5C**) and width (**Fig. S5E**). The R303E K304E mutation did not reduce induction of 1° cell fate in the VPC system (**Fig. 5E**; **S5F**), suggesting that LET-60/Ras does not activate AGE-1 in VPC fate patterning. The R303E K304E mutation also did not alter lifespan (**Fig. S5G,H**), as might be expected for the likely general reduction-of-function conferred by the *age-1(ag12)* mutation, consistent with *ag12* more generally destabilizing the AGE-1 protein. Thus, we hypothesize that *age-1(re353re377*rf*[age-1::mNG::2xHA R303E,K304E])* specifically targets the binding interface of LET-60/Ras and the RBD of AGE-1/PI3Kact, thus revealing a novel phenotype and hence regulatory control of animal growth.

### The T35S putative Raf-selective mutation in LET-60/Ras phenocopies the putative RBD-deficient mutation in AGE-1/PI3Kcat

To further test whether the R303E K304E mutation abrogates LET-60/Ras binding to AGE-1/PI3Kact, we selectively mutated the effector binding loop (EBL) of endogenous LET-60/Ras to reduce binding to AGE-1/PI3Kact. A series of effector binding mutations were previously defined via yeast two-hybrid analysis and cell-based studies that selectively supported activation of certain proto-oncogenic effectors by constitutively activated Ras (Rodriguez-Viciana et al., 1997; White et al., 1995; Wolthuis & Bos, 1999). The efficacy of this mutation was validated via structural biology and *in vitro* binding (Pacold et al., 2000). Of the canonical EBL mutants, T35S is Raf-selective, E37G is RalGEF-selective, and Y40C is PI3Kcat-selective. However, these mutations do not necessarily fully cleanly or completely interfere with function: they may not retain 100% activation of effectors nor specifically block one effector.

We previously used constitutively activated LET-60 G12V,T35S and G12V,E37G via VPC-specific transgenesis; both functioned *in vivo* as expected from the mammalian literature: G12V,T35S appeared to activate the LET-60/Ras>LIN-45/Raf 1°-promoting pathway and G12V,E37G appeared to activate the LET-60/Ras>RGL-1/RalGEF 2°-promoting pathway (Zand et al., 2011). However, LET-60/Ras activation of LIN-45/Raf is essential for viability in *C. elegans* (Yochem, Sundaram, & Han, 1997). Consequently, we cannot use other EBL mutations in endogenous LET-60/Ras because of the expectation they will eliminate activation of LIN-45/Raf and hence result in lethality. Thus, we are constrained to using the T35S mutation in endogenous LET-60/Ras to abrogate LET-60 binding to AGE-1. This mutation is also expected to abrogate RalGEF signaling, as well as perhaps other, non-proto-oncogenic effectors.

We used CRISPR/Cas9-dependent genome editing to generate *let-60*(*re378[T35S]*), which encodes T35S mutant LET-60/Ras. T35S mutant animals were superficially wild-type in every regard except for increased stature, similar to the R303E K304E RBD mutant of AGE-1 (**Fig. 5D**; **S5F**). Unfortunately, we cannot generate the G12V T35S constitutively active LET-60/Ras because the G12V activating mutation in *C. elegans* confers Raf-dependent lethality (J. Mardick and D. Reiner, unpublished). We propose that a LET-60/Ras>AGE-1/PI3Kcat imposes restriction of animal stature, such that reduced signal enables increased animal size and increased signal might decrease animal size.

### Mutations in the binding interface of LET-60/Ras>AGE-1/PI3Kcat reveal an unexpected role in the dauer diapause decision

We used the T35S mutation in LET-60/Ras and the R303E K304E mutation in the RBD of AGE-1/PI3Kcat to evaluate the role of LET-60/Raf>AGE-1/PI3Kcat signaling in dauer formation. Unexpectedly, we observed that under multiple conditions both T35S and R303E K304E mutants reduced dauer formation (**Fig. 6A-C**). Consequently, we propose that the LET-60>AGE-1 signal promotes formation of dauers.

**Figure 6.**
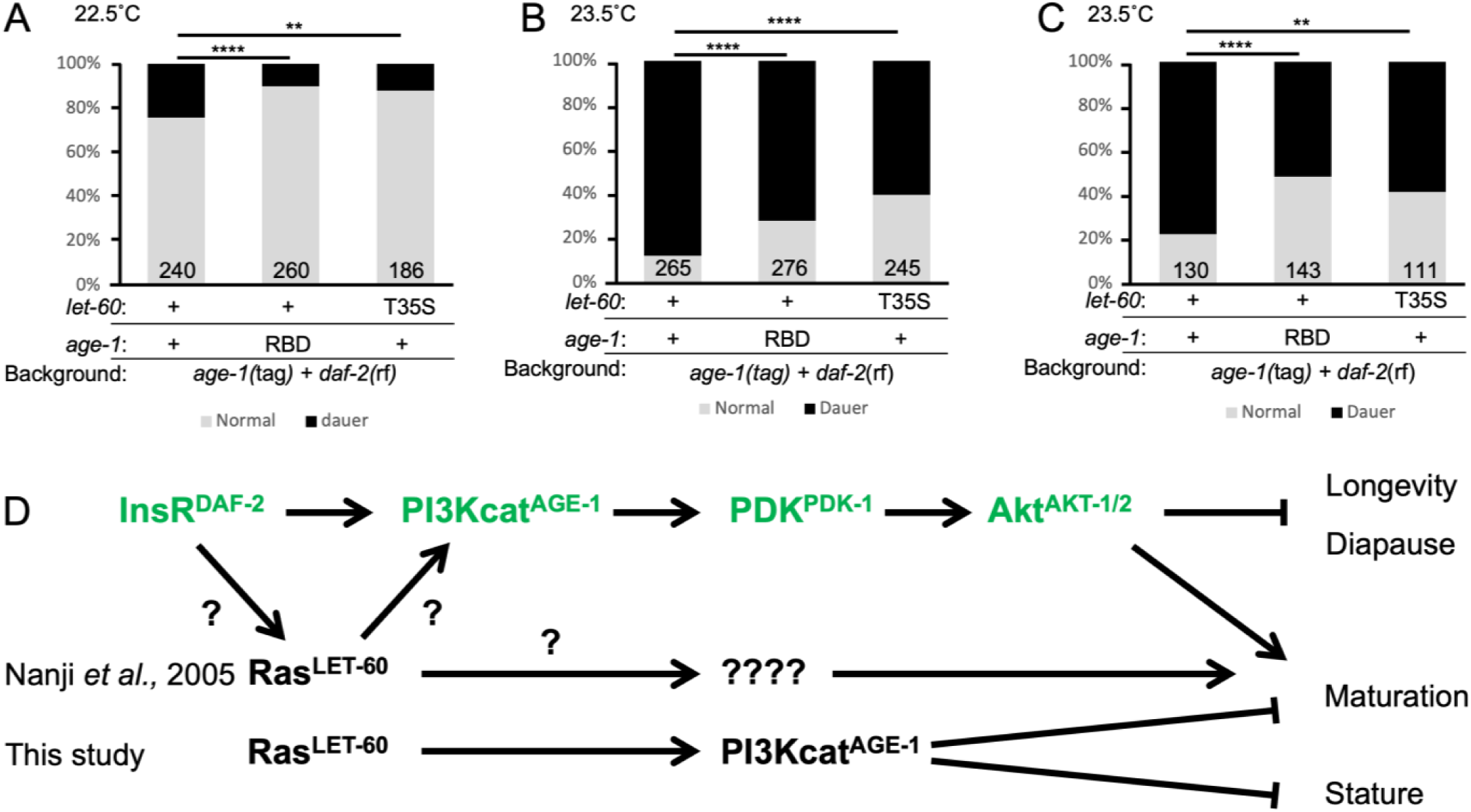
Putative defective association of LET-60/Ras with the AGE-1 RBD reduces dauer formation. **(A-C)** Percent dauer formation of *daf-2(m577*rf) animals at different temperatures with *age-1(tag)* alone, RBD mutation or *let-60(T35S)* putative PI3K defective mutation. (The weaker *daf-2(m577)* was used because *daf-2(e1370*) conferred 100% dauer formation at these temperatures.) Animals in the same graph were scored concurrently. P values were calculated by Pearson‘s chi-squared test. For p values, * < 0.05, ** < 0.01, *** < 0.001, **** 0.0001. **(D)** A schematic of the dauer regulatory network, determined by a combination of published results and the novel interactions shown in this study. The core IIS pathway, DAF-2/InsR>AGE-1/PI3Kcat>PDK-1/PDK>AKT-1/2/Akt, is as found in *C. elegans* and other systems. Upstream sensory inputs converge on IIS. The pathway inhibits dauer diapause and promotes developmental maturation and reproduction. Prior experiments (Nanji *et al*., 2005) with G13E constitutively activating LET-60/Ras suppressed dauer formation caused by reduction-of-function DAF-2/InsR, arguing that LET-60/Ras promotes reproductive fate at the expense of dauer diapause. Our results with disruption of the LET-60/Ras>AGE-1/PI3K biding interface reduced dauer formation caused by reduction-of-function DAF-2/InsR, suggesting LET-60/Ras>AGE-1/PI3K inhibits maturation/reproductive development and promotes dauer diapause. This contradiction could be reconciled by different activities in distinct tissues, or the constitutive activation of all LET-60 activities, including potentially effectors LIN-45/Raf, RGL-1/RalGEF and other non-proto-oncogenic effectors, cause different outcomes with different effectors.

IIS components in *C. elegans* were identified largely through roles in control of the dauer diapause decision, resulting in a model of a DAF-2/InsR>AGE-1/PI3Kcat>PDK-1/PDK>AKT-1/2 cascade repressing dauer development and promoting reproductive development (**Fig. 6D**; Fielenbach & Antebi, 2008). One study showed that the *let-60(n1046*gf) G13E moderately activating mutation suppressed reduced function mutations in DAF-2/InsR that precociously promote dauer formation (dauer constitutive, or Daf-c; Nanji, Hopper, & Gems, 2005). Moreover, reduction of LET-60/Ras function was predicted to weakly promote formation of dauers. Together, these results led to the model that LET-60/Ras signaling functioned downstream of DAF-2/InsR to repress dauer formation and promote reproductive development. This signal was speculated to be through AGE-1/PI3Kact but that hypothesis could not be tested.

Our results contradict that finding. Predicted reduction of RBD function and reduction of LET-60/Ras activation of AGE-1/PI3Kcat binding of LET-60/Ras both reduce dauer formation. Consequently, we hypothesize that LET-60>AGE-1 inhibit developmental progression to maturation at the expense of dauer formation, i.e. promote dauer diapause (**Fig. 6D**).

A major methodological difference is that the previous mutations either reduced or increased all signaling functions of LET-60/Ras. Perhaps these genetic perturbations alter multiple functions of LET-60, at least one of which strongly promotes reproductive development, whereas our missense mutations more selectively alter LET-60 and AGE-1 functions. Alternatively, perhaps LET-60 acts at multiple positions in the dauer regulatory network, or in multiple tissues in the animal. Some of these could be positive and some negative signals, and some via AGE-1/PI3Kcat and others through another effector, like LIN-45/Raf. Regardless, these fundings illustrate the utility of using selective mutations driven by structural knowledge when interrogating biological functions *in vivo*.

## DISCUSSION

The goal of this study was to use structural insights to guide *in vivo* editing of the sole Class IA PI3K catalytic subunit in *C. elegans*, AGE-1, and to define organismal consequences of discrete mutations. By combining structural threading with CRISPR/Cas9-dependent genome editing, we introduced a functional C-terminal fluorescent tag, an activating mutation modeled on oncogenic PI3Kcat/p110α alleles, and a Ras-binding–deficient mutation. These tools allowed us to probe AGE-1 function in lifespan, vulval development, growth, and dauer entry.

Our tagging strategy highlights the value of structure-based design for engineering large multidomain proteins. By introducing an extended flexible linker between AGE-1 and the fluorescent protein, we avoided steric clashes predicted from crystallographic studies and supported by AlphaFold3 models. The tagged protein retained full function in lifespan and vulval assays, including in sensitized *let-60(*gf) and *daf-2(*rf) genetic backgrounds, allowing accurate visualization of endogenous expression. This tagged protein now provides a versatile platform for future structure-function studies and biochemical analyses. Likewise, structural modeling of the helical and Ras-binding domains enabled us to design and generate alleles that either phenocopied known oncogenic mutations or selectively disrupted Ras interaction without destabilizing the protein. These results underscore the power structural information to produce design precise, interpretable alleles for whole-animal studies.

### Animal stature

An unexpected finding was the increase in body size in both Ras-binding-deficient AGE-1 mutants and the complementary LET-60 effector-binding mutant. To our knowledge, no parallel has been described for PI3K in other systems. In mammalian cells, PI3K activation generally promotes growth. Yet in *C. elegans* our results suggest that Ras>PI3K signaling restrains growth. This points to a non-canonical, possibly tissue-specific role for LET-60>AGE-1. Determining the site of action – whether neuronal, intestinal, or hypodermal – will require conditional or tissue-specific genetic manipulations.

In *Drosophila*, InsR signaling through mTORC1 is a major positive regulator of cell and body size (Oldham & Hafen, 2003). Similarly, *C. elegans* grown on nutrient-rich HB101 *E. coli* bacteria are 1.6-fold larger than animals grown on standard OP50, and this effect is partially reversed in DAF-2/InsR mutants (So, Miyahara, & Ohshima, 2011). Moreover, some *daf-2* reduction-of-function alleles produce longer animals (McCulloch & Gems, 2003). Together, these observations suggest that DAF-2/InsR exerts pleiotropic control overgrowth, lifespan, and the decision between diapause vs. reproductive development, with context-dependent effects on body size. Within this broader framework, our selective mutations in LET-60 and AGE-1 reveal a specific for LET-60>AGE-1 signaling in restraining growth, opening new avenues for dissecting how these conserved pathways shape organismal size.

Our dauer assays likewise revealed unexpected complexity. Previous models positioned LET-60 downstream of DAF-2/InsR, promoting reproductive development and repressing dauer entry (Nanji et al., 2005). In contrast, both Ras binding-deficient AGE-1 and AGE-1 binding-defective LET-60 alleles reduced dauer formation, suggesting that LET-60>AGE-1 signaling can promote dauer entry. These results raise the possibility of multiple LET-60>AGE-1 inputs, acting in different tissues or developmental windows, with opposing effects on dauer regulation. Dauer entry is further influenced by parallel pathways, including the EAK (<u>e</u>nhancer of *<u>ak</u>t-1*) genes (Williams, Dumas, & Hu, 2010), whose relationship to Ras>PI3K signaling remains unresolved. Targeted manipulations with cell type-specific alleles will be needed to clarify these interactions.

A critical advantage of our approach is the use of CRISPR/Cas9-dependent genome editing to introduce precise lesions with minimal off-target effects. The resulting strains carry orders of magnitude fewer background mutations than strains derived via chemical mutagenesis or natural isolates (Denver et al., 2009; Sarin et al., 2010).

In summary, our structure-based engineering of AGE-1 demonstrates how rationally designed mutations can illuminate domain-specific functions *in vivo*. The activating E630K mutation confirmed conserved roles of PI3K in lifespan and growth factor signaling. Selective disruption of Ras-PI3K binding revealed novel roles in growth restraint and dauer regulation. Together, these findings show how integrating structural models with the genetic tractability of *C. elegans* can unveil unexpected biology and provide new entry points into conserved signaling pathways.

## METHODS

### *C. elegans* handling and genetics

Strains used in this study were derived from the N2 Bristol wild type and grown at 20°C on NGM plates seeded with *E. coli* strain OP50. *C. elegans* strains used are listed in Table S1, Oligonucleotides in Table S2, CRISPR reagents in Table S3. Nomenclature is as described (Tuli, Daul, & Schedl, 2018).

### CRISPR/Cas9-dependent genome editing

Guide RNAs were selected using a combination of three criteria. First, we prioritized sequence features shown to improve Cas9 activity: where possible, guanine (G) rather than thymine (T) nucleotides were chosen at positions -1/-2/-4, or the GCGG sequence in preference over Ts at positions −1 through −4 (Farboud & Meyer, 2015; Wang, Wei, Sabatini, & Lander, 2014). Second, predicted specificity and efficiency were evaluated using the CRISPOR algorithm (https://crispor.gi.ucsc.edu/; (Concordet & Haeussler, 2018), which incorporates the original MIT specificity score. Third, predicted efficiency was further assessed using the WU-CRISPR algorithm (http://crisprdb.org/wu-crispr/; (Wong, Liu, & Wang, 2015)).

Injection mixes were formulated as described (Fakieh & Reiner, 2025; Ghanta, Ishidate, & Mello, 2021). Specifically, we used final concentrations of 0.25 mg/ml SpCas9 (PNABio), 0.1 mg/ml universal tracrRNA, 0.028 mg/ml crRNAs for the *dpy-10* co-CRISPR marker and the edit of interest, repair templates for the dominant *dpy-10(cn64*gf) co-CRISPR marker and the repair template of interest to 3.3 mM (Table S3). Mixes were assembled in an RNAse-free bench space and incubated at 37°C for 15 min.

F1 animals expressing the *dpy-10(cn64*gf) Rol co-CRISPR marker (Arribere et al., 2014) were picked singly to plates. After F2 embryos were laid, F1s were picked into tubes – sometimes singly and sometimes in pools of 2 – and lysed for single-worm PCR. Non-Rol or -Dpy F2 animals from PCR positives were PCR amplified for homozygous edits. The edited region from resulting homozygotes was amplified and Sanger sequenced.

### Imaging

Live animals were mounted on 3% agar pads with 5 µl of 2 mg/ml tetramisole in M9 buffer. Differential Interference Contrast (DIC) images were captured using the Nikon 224 Eclipse Ni Microscope with 60x oil objective. Confocal fluorescent micrographs were captured using a Ti2-Nikon inverted microscope equipped with a Yokogawa CSU-W1 Spinning Disk and a Photometrics Prime BSI camera. Green fluorescence (mNeonGreen) was captured at 488 nm. NIS Elements Version 4.30 software was used to capture and process images.

### Quantifying animal phenotypes

Synchronized embryos were obtained by picking adults (24 hours post-late L4) onto plates with food and picking off 1 hour later.

Animal size was quantified using synchronized adults 72 hours after egg laying. DIC photos were taken using a spinning disk confocal microscope with the 40x objective. Length and width of animals were analyzed using NIS Elements Version 4.30 software. Length was measured as distance from the anterior to posterior (A-P) end and width as the ventral to dorsal (V-D) at the A-P midpoint where the vulva is located.

For lifespan assays, 20 L4 animals were picked to each plate, multiple plates for each genotype. Animals were transferred and assessed every 24 hours after initial picking until production of embryos ceased. Total number of surviving animals was quantitated every other day.

Strains for dauer assays were propagated at 15°C. Ten young but gravid adults were picked to each plate for egg laying at the temperature indicated and picked off for approximate synchrony of embryos. Dauer vs. adult animals for each plate were quantitated 72 hours later.

The role of AGE-1/PI3Kcat signaling in patterning of cell fate of vulval precursor cells (VPCs) fate was scored as described (Fakieh & Reiner, 2025; Shin et al., 2018). Briefly, we counted ectopic 1° VPCs induced by gain of function of the *C. elegans* ortholog of Ras, *let-60(n1046*gf), with and without edits in *age-1*. Late L4 animals were mounted and imaged via DIC (see above). Ectopic 1°s were identified as single-lobed invaginations compared to the three-lobed invagination of the normal vulva. In strains harboring *let-60(n1046*gf), we have observed drift of the strong of the phenotype of ectopic 1° pseudovulvae, typically becoming more severe. As previously described, we carefully avoided multi-generational plate culture of animals, either analyzing freshly from a thawed strain, recently isolated as a homozygote from a cross, or recovered from a starved, parafilmed plate that was stored immediately after thawing. Strains not fitting the established baseline range of 1.2-1.5 ectopic 1° pseudovulvae are discarded and the strains restarted as described (Zand et al., 2011).

### Detection of tagged endogenous protein

For immunoblotting, mixed stage animals were washed from plates using M9 buffer and lysed in 4% SDS loading buffer by boiling at 90°C for 5 minutes. Lysates were run on 4%–15% SDS gel (Bio-Rad) and blotted on Immobilon-P Membrane, PVDF (EMD Millipore, IPVH00010). Anti-HA antibody (Proteintech 51064-2-AP) and anti-α-tubulin antibody (Sigma-Aldrich T6199) were diluted 1:2000 in blocking buffer (6% w/v non-fat dry milk in PBST). HRP-conjugated goat anti-mouse secondary antibody (MilliporeSigma 12-349) was diluted in 1:5000 in blocking buffer. Chemiluminescent detection was performed using ECL reaction (Thermo Fisher Scientific), and detected using the Bio-Rad Chemidoc MP Imaging System Hood III.

### Protein structural analysis and model construction

Structure threading used PHYRE2.2 (Powell, Islam, David, & Sternberg, 2025) to obtain a model of the AGE-1+AAP-1 structure, which was then used to 3D align with earlier PI3KCA/p85 structures. The sequence of LET-60 was threaded onto the structure of MRAS bound to PI3KCA (Czyzyk et al., 2025). Structures were visualized with PyMol 2.6 (Schrödinger, LLC) and earlier versions (https://www.pymol.org/). Protein Data Bank accession numbers used were 4OVU, 4OVV, 9B4T (Burley et al., 2025).

### Software and statistical/data analysis

Data analysis and graphing was performed using Prism 9 (version 9.0.1) and Microsoft Excel (version 16.45). Ageing assay data analysis and the Kaplan-Meier graph were used the online survival analyzing tool on https://sbi.postech.ac.kr/oasis/ (Yang et al., 2011). Protein alignments used Clustal Omega, domain analysis used SMART and Prosite.

## ACKNOWLEDGEMENTS

We thank Deverie Bongard, Fred Ausubel and Gary Ruvkun (Harvard Medical School) for searching for the frozen aliquot of the strain harboring *age-1(ag12)* during the COVID-19 pandemic, Andy Golden (NIDDK) for general generosity and sharing his composite CRISPR protocol, Robyn Tanguay (Oregon State U) for introducing the phrase “model discipline” to D. Reiner, L. Vergara of the Center for Advanced Imaging for technical advice, and members of the Reiner lab for helpful discussions. Some strains were provided by the *Caenorhabditis* Genetics Center, which is funded by the NIH Office of Research Infrastructure Programs (P40 OD010440). Wormbase was used routinely (Sternberg et al., 2024). This work was supported by NIH grants R35GM144237 and R03CA289854 to DJR.

## Contributions

conception, design, acquisition of data, analysis and interpretation of data: YW, DR; drafting and revising the manuscript: YW, DR; critical technical mentoring and/or generation of key reagents: TD, NR; structural analysis: KR, YW, DR.

## Ethics

All ethics and safety guidelines were observed. There are not conflicts of interest to report.

## Data availability

All data are presented in the manuscript; no large data sets requiring upload were generated. All reagents are available upon request or are being sent to key repositories, *e.g.* CGC, Addgene.org.

---

**Figure S1.**
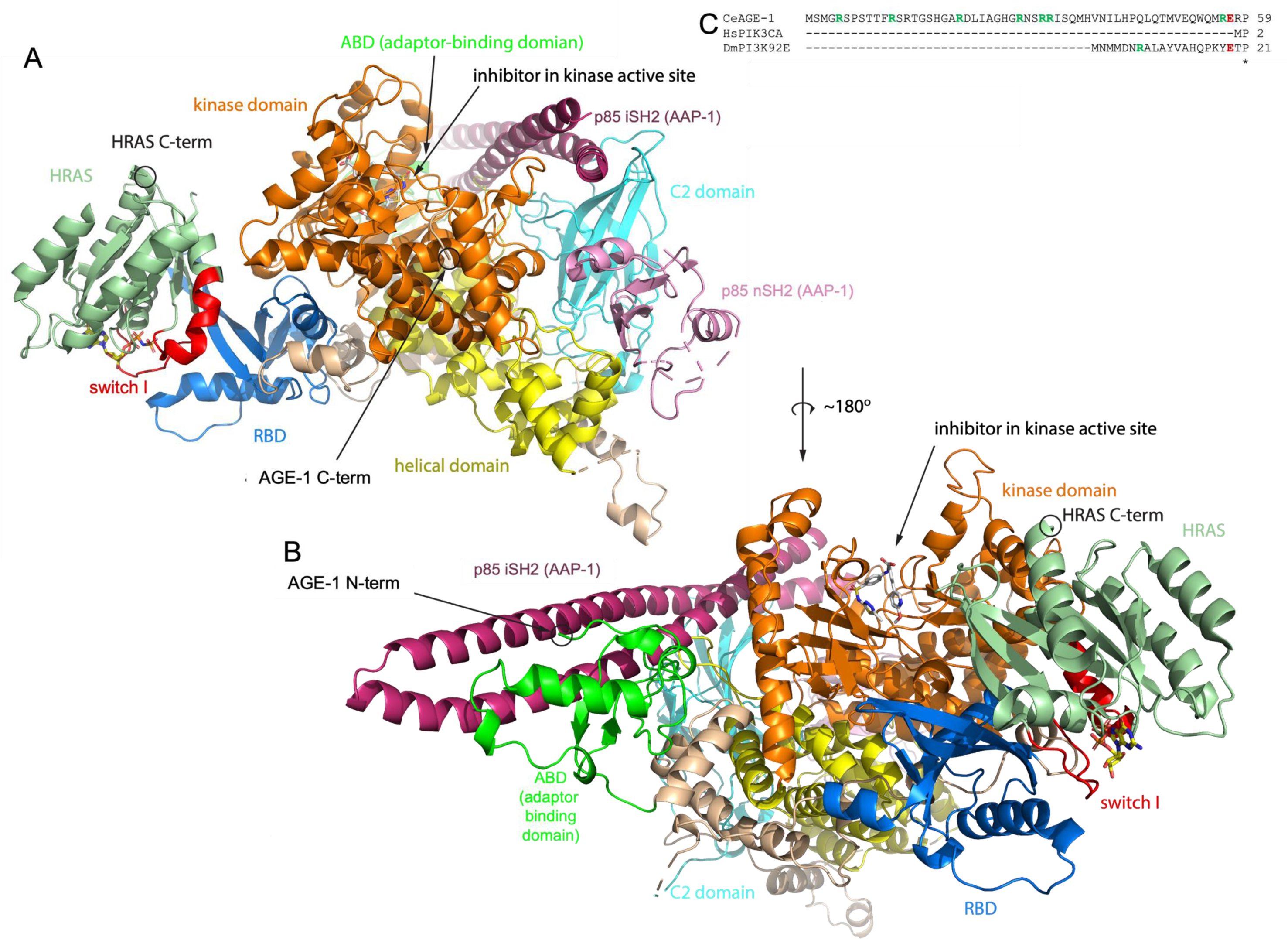
Structural model of human HRAS, PI 3-Kinase catalytic alpha, and PIK3R1 p85. **(A)** A *pre hoc* model of *C. elegans* AGE-1+AAP-1 threaded onto PI3Kcat alpha and p85 with an HRAS structure superimposed. HRAS is bound to non-hydrolysable GTP analog GMPPNP, is colored in pale green, and the Switch II region that binds effectors and shifts upon GTP binding is colored in red. This model of threaded AGE-1+AAP-1 bound to HRAS specifically led us to tag the endogenous AGE-1 protein at the C-terminus. The plasma membrane in this model is located above the enzyme, as indicated by the C-terminus (indicated) of HRAS, which is directed to the PM. The C-terminus is predicted to be sufficiently far from the PM to avoid steric interference of an ∼27 kDa fluorescent protein, with a 30 residue GASx10 linker sequence added to decrease risk of interference. The N-terminal SH2 domain (nSH2) is shown partially structured. The C-terminal SH2 domain (cSH2) is not shown, as is typical, as are the domains of p85 missing in p50 and p55. **(B**) A 180° rotation of the structural model reveals the predicted location of the N-terminus. Tagging with an FP might interfere with general PM association. **(C)** The unstructured N-terminal extensions of *C. elegans* AGE-1 and *Drosophila* PI3K92E are shown, with Arginine residues bolded and in green. We hypothesize that the basic N-terminus of AGE-1 may provide an electrostatic charge with the acid PM, which might be destabilized by an FP tag, another reason to avoid the N-terminus of AGE-1 for tagging.

**Figure S2.**
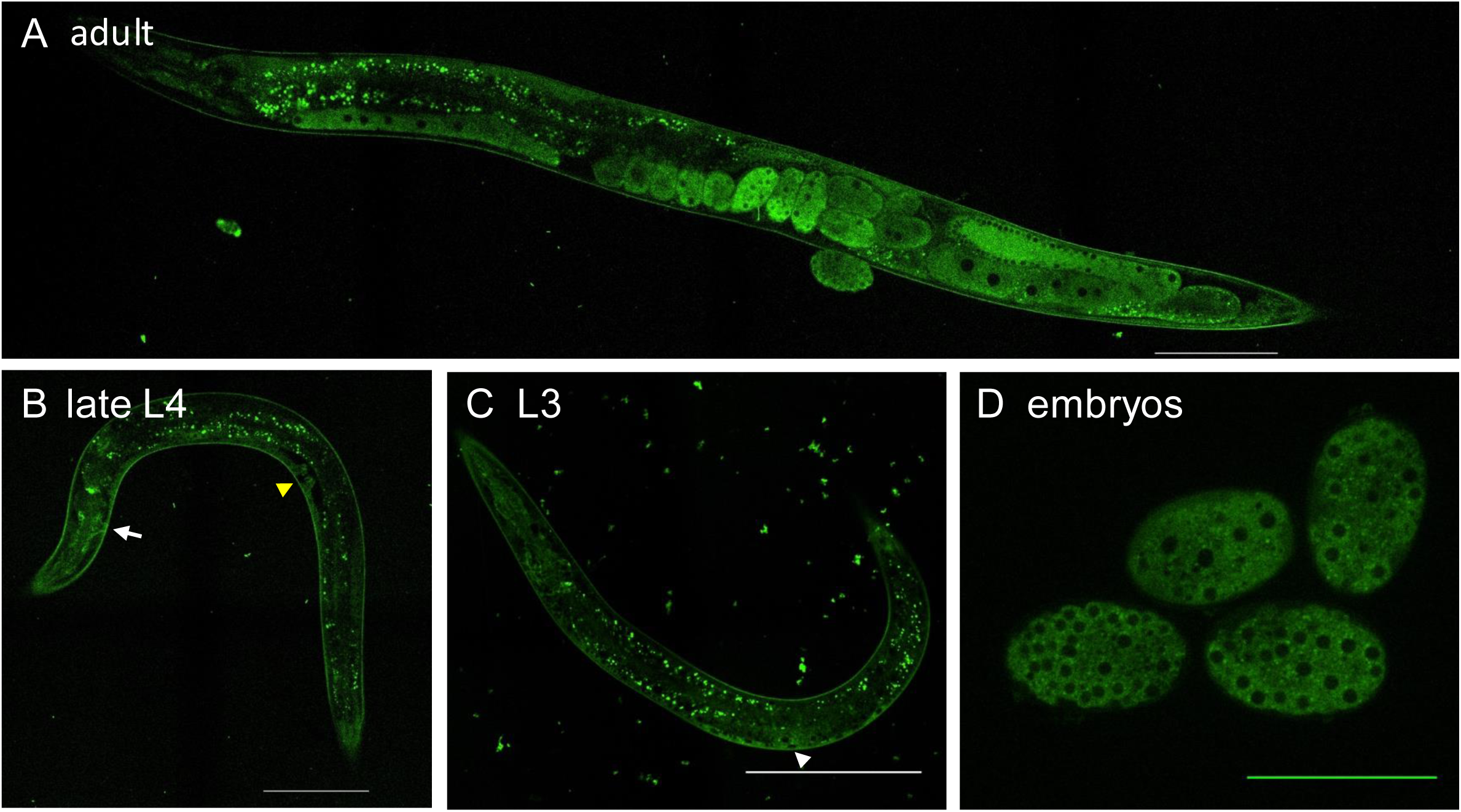
Spinning disk confocal photomicrographs (488 nm) of *age-1(re353[age-1::linker::mNG::2xHA])* animals. **(A)** Adult. **(B)** Late fourth larval stage (L4). White arrow = nerve ring neuropil (a bundle of many neurites) and perhaps the excretory duct/pore. White arrow = nerve ring neuropil (a bundle of many neurites.) Yellow arrowhead = morphogenetic vulva. **(G)** Third larval stage. White arrowhead = P6.p of the VPCs, flanked by other VPCs along the ventral surface. See Figure 2 legend: bright green spots are autofluorescence from intestinal lipid droplets. Dark circles are nuclei. **(H)** Embryos. We consistently observe speckles in embryos, presumably cytosolic, but do not know the source. White scale bars = 100 µm. Green scale bar = 50 µm.

**Figure S3.**
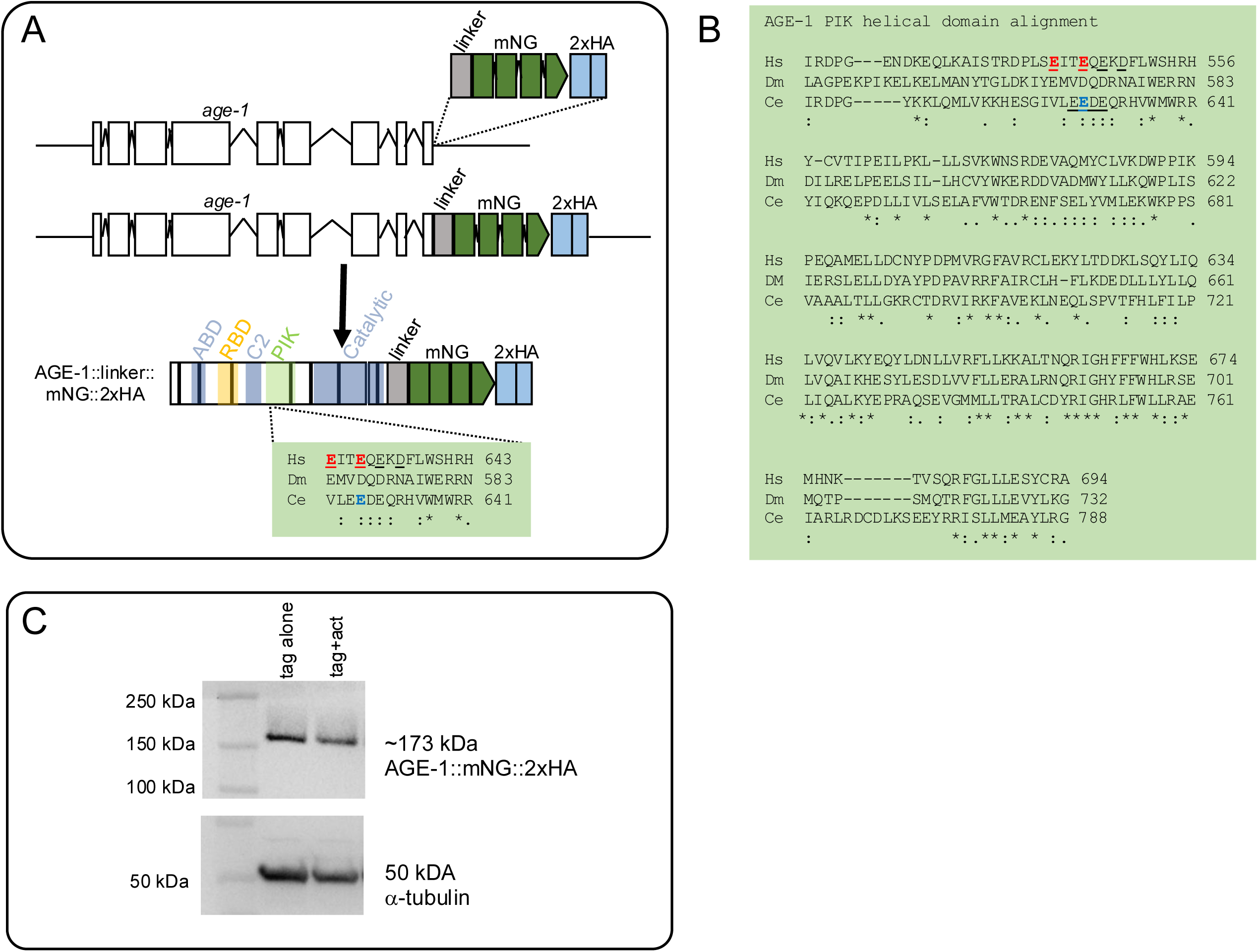
Support for the gain-of-function mutation in AGE-1. **(A)** Schematic of the edited *age-1* gene and protein with tag (see Fig. 2A) or with the E630K mutation. **(B)** An alignment of the entire PIK/helical domains. Hs = *Homo sapiens*, Dm = *Drosophila melanogaster,* Ce = *Caenorhabditis elegans*. Human E542 and E545 residues altered to K in oncogenic lesions in the PIK/helical domain are shown in red, the nematode E630 residue altered to to K in the gain-of-function is shown in blue. Each protein has four acidic residues in this region, just clustered in nematodes and more diffuse in humans. While the helices in the predicted structure are of the PIK/helical domain reasonably well conserved, the primary sequence is not. **(C)** Immunoblotting with anti-HA indicates the E630K mutation did not alter protein stability of expression. The western shown is the same shown for sizing in Fig. 2B, from which the E630K lane and the RBD lanes (see **Fig. S5C**, below) were cropped.

**Figure S4.**
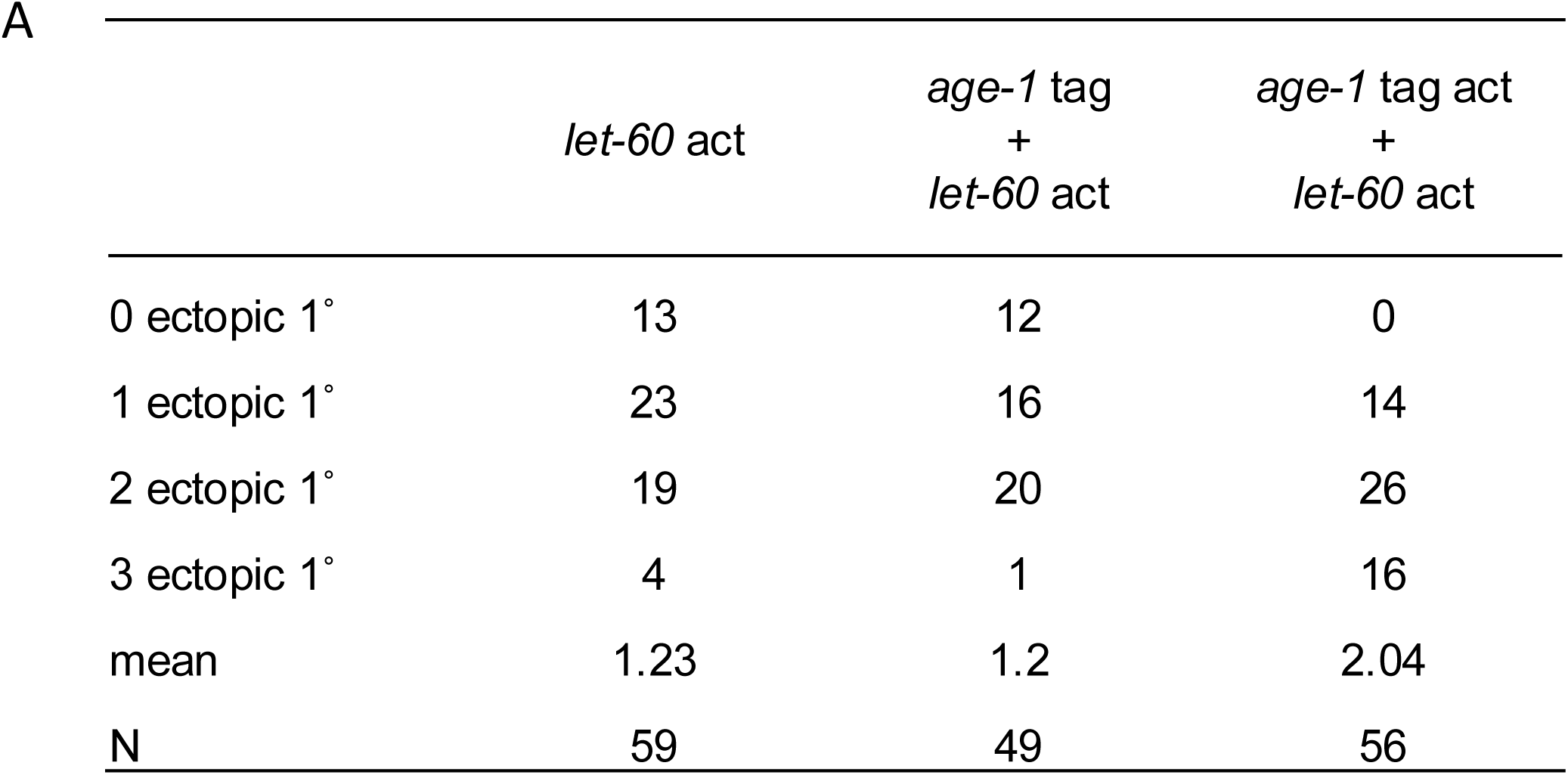
Raw data for Figure 4F. All animals were scored concurrently.

**Figure S5.**
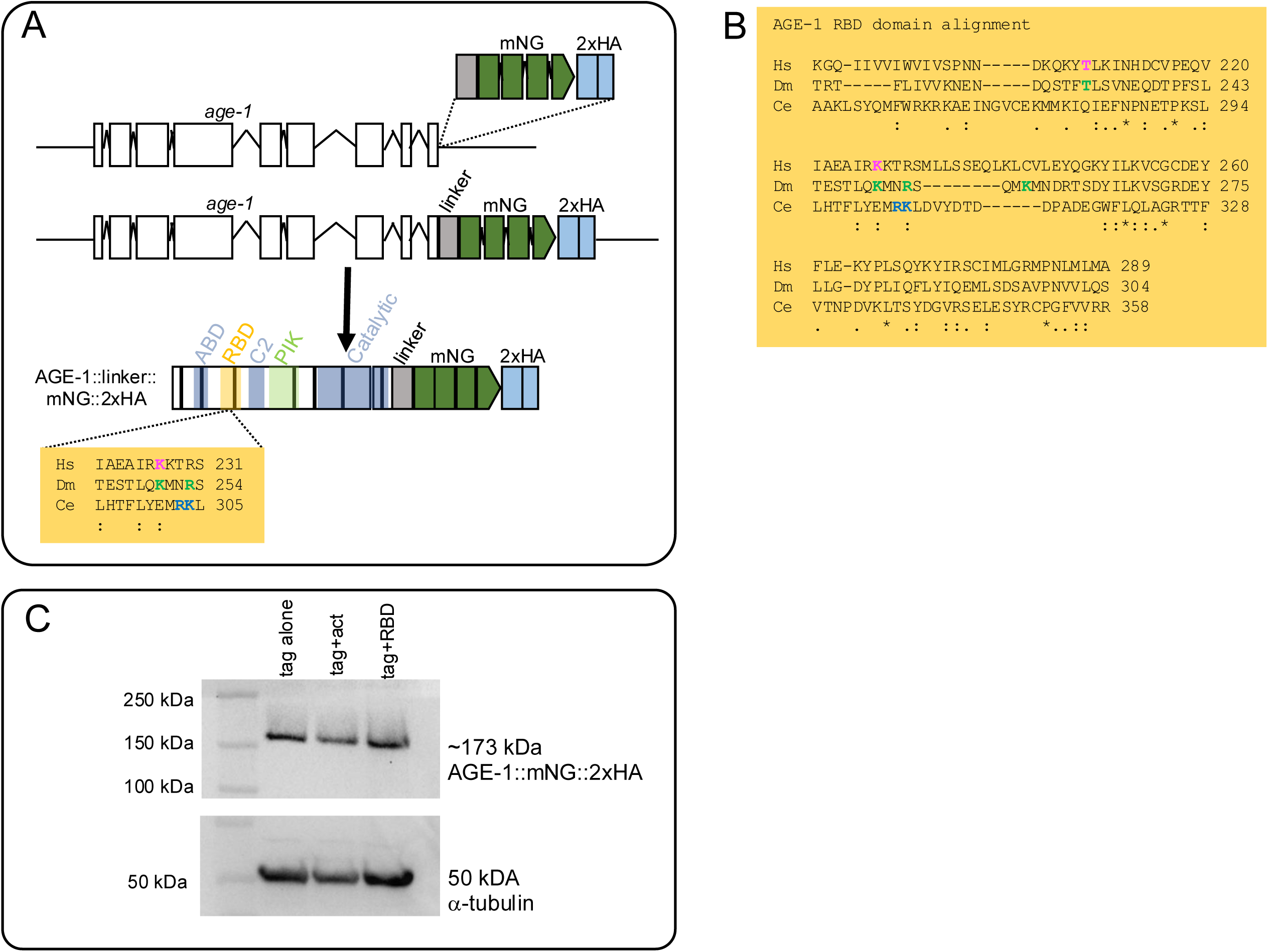
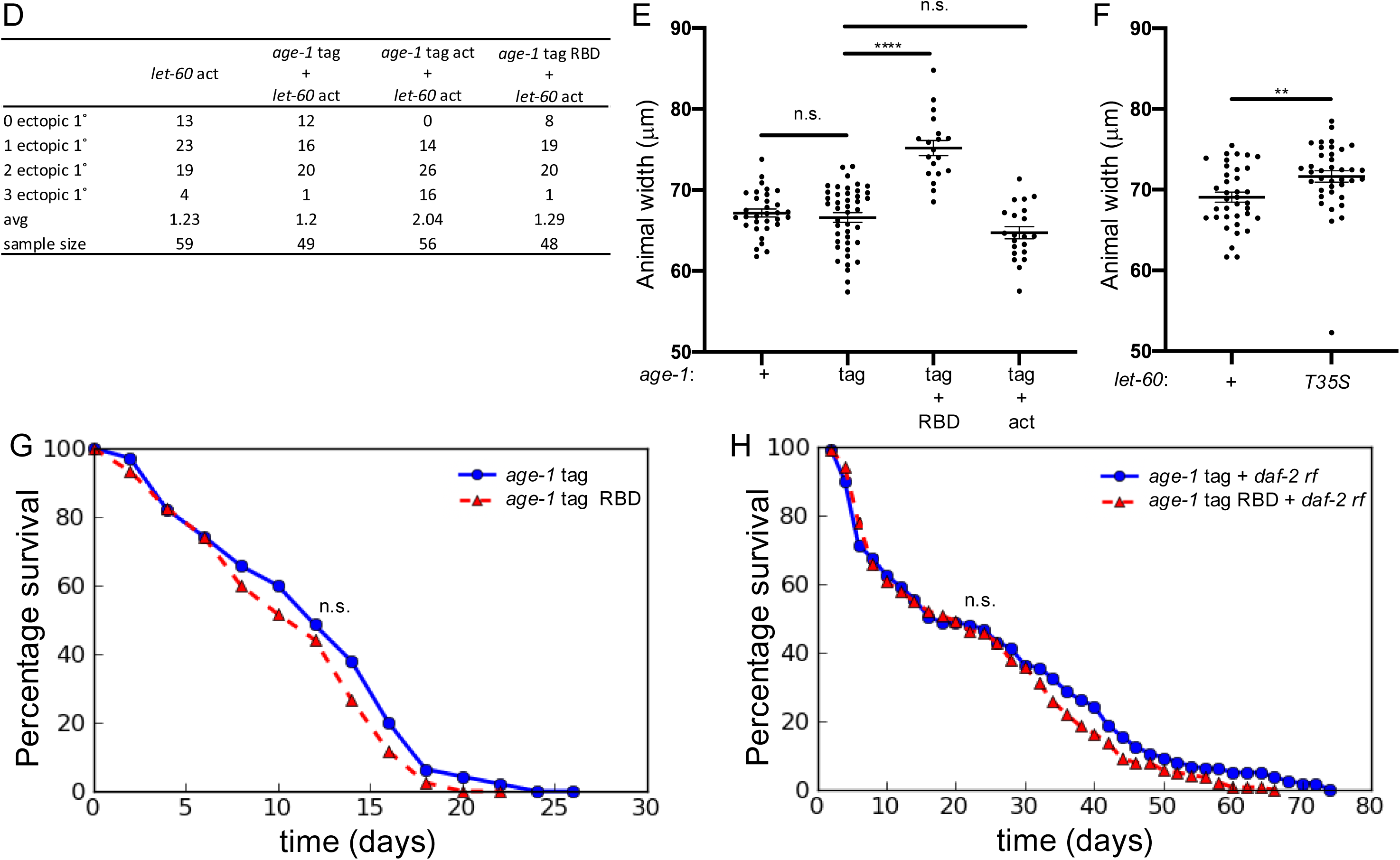
Support for the RBD in AGE-1/PI3Kcat. **(A)** Schematic of the edited *age-1* gene and protein with tag (see Fig. 2A**, S3A**) or with the R303E K304E mutation. **(B)** An alignment of the entire RBD domains. Hs = *Homo sapiens*, Dm = *Drosophila melanogaster,* Ce = *Caenorhabditis elegans*. Human T208 and K227 residues altered to D and A, respectively, in the human RBD deficient mutant (Gupta *et al*. 2007) are shown in pink. *Drosophila* T231, K250, R253 and K257 altered to D, A, A, and A, respectively in the *Drosophila* RBD deficient mutant (Orme *et al*, 2006) are shown in green. With the relatively poor conservation of the RBD, we did not mutate these specifically but rather mutated structure-based residues. The nematode R303 K304 residues altered to to E in the RBD reduction-of-function mutant are shown in blue. **(C)** Immunoblotting with anti-HA indicates the R303E K304E mutations did not alter protein stability of expression. The western shown is the same shown for sizing in Fig. 2B, where the E630K lane and the RBD lanes were cropped. **(D)** Data for VPC scoring in Fig. 5E. **(E)** Animal width in microns as measured from DIC photomicrographs for RBD and act mutations in AGE-1 as shown in Fig. 5C. **(F)** Animal width in microns as measured from DIC photomicrographs for the LET-60 T35S putative Raf-selective mutant vs. wild type as shown in as shown in Fig. 5D. **(G)** Kaplan-Meier curve for lifespan of tag vs tag+RBD mutants. N= 140 for *age-1* tag, N = 120 for *age-1* tag+RBD. **(H)** Kaplan-Meier curve for lifespan of tag vs tag+RBD mutants in the *daf-2(e1370rf)* reduced function mutant background. N= 160 for *age-1* tag + *daf-2(e1370*rf); N = 120 for *age-1* tag+RBD + *daf-2(e1370*rf). P value for animal length and width was calculated using a *t* test, and animal aging was calculated using Log-rank (Mantel-Cox) test and ANOVA. n.s. = not significant, ** ≤ 0.01, **** ≤ 0.0001.

**Table S1.** Strain list.

| Strain | Genotype | Impact |
| --- | --- | --- |
| DV4018 | *age-1(re353tag[age-1::mNG::2xHA])* II | FP tag |
| DV4070 | *age-1(re353re377rf[age-1::mNG::2xHA(R303E,K304E)])* II | RBD |
| DV4138 | *age-1(re353re392gf[E630K])* II | Activated |
| DV4019 | *age-1(re353tag)* II; *daf-2(m577rf)* III |  |
| GR1318 | *pdk-1(mg142gf)* X | Activated |
| DV4179 | *age-1(re353re392gf)* II; *daf-2(e1370rf)* III |  |
| DV4180 | *age-1(re353tag)* II; *daf-2(e1370rf)* III |  |
| DV4181 | *age-1(re353re377rf)* II; *daf-2(e1370rf)* III |  |
| DV4019 | *age-1(re353tag)* II; *daf-2(m577rf)* III |  |
| DV4075 | *age-1(re353re377rf)* II; *daf-2(m577rf)* III |  |
| DV4139 | *age-1(re353re392gf)* II; *daf-2(m577rf)* III |  |
| DV4094 | *let-60(re378 [T35S])* IV |  |
| DV4112 | *daf-2(m577rf)* III; *let-60(re378 [T35S])* IV |  |
| DV4300 | *age-1(re353tag)* II; *let-60(n1046gf)* IV |  |
| DV4301 | *age-1(re353re392gf)* II; *let-60(n1046gf)* IV |  |
| DV4302 | *age-1(re353re377rf)* II; *let-60(n1046gf)* IV |  |

**Table 2.**
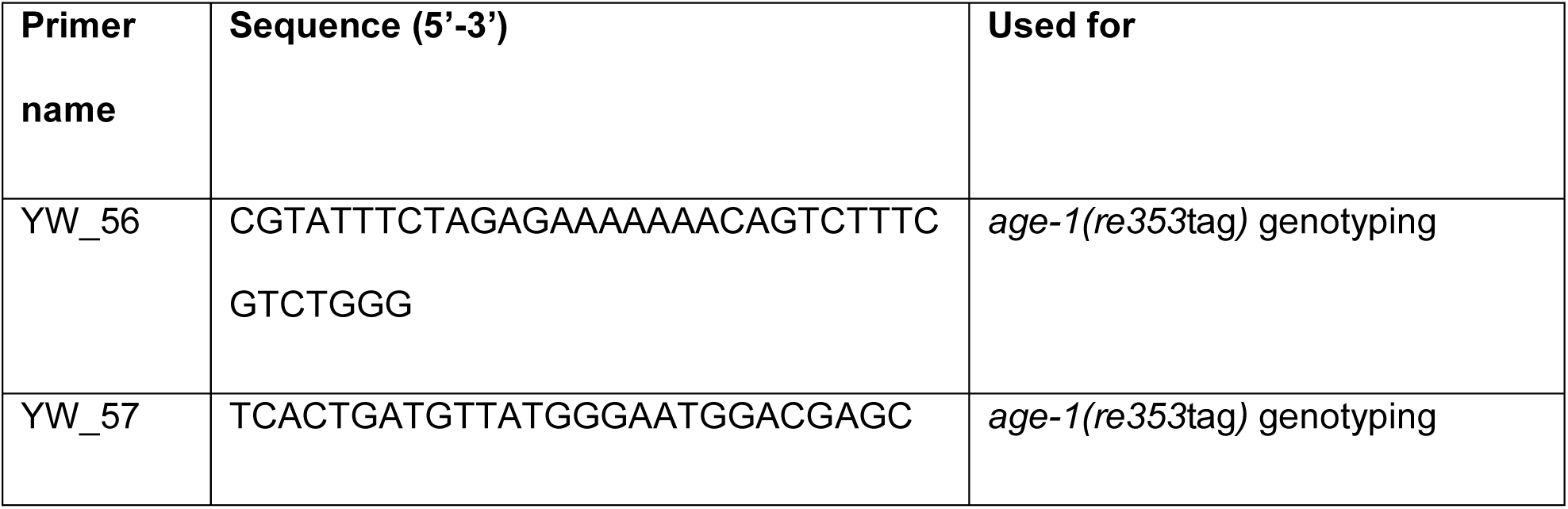
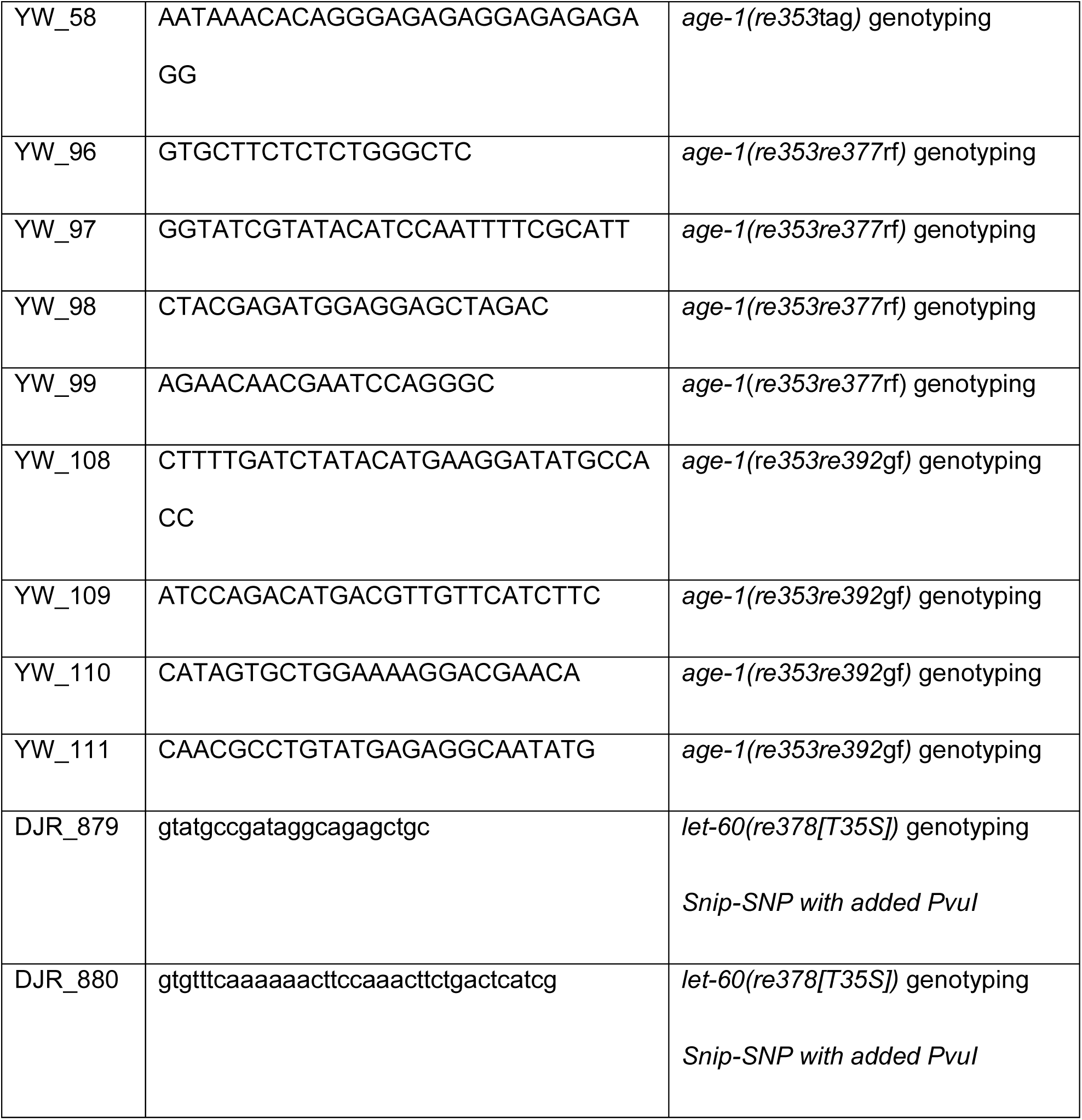
Primer list.

**Table 3.**
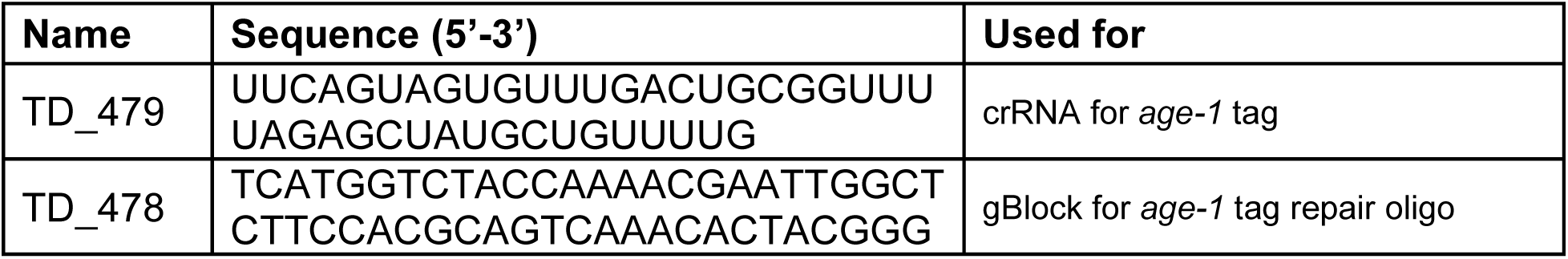
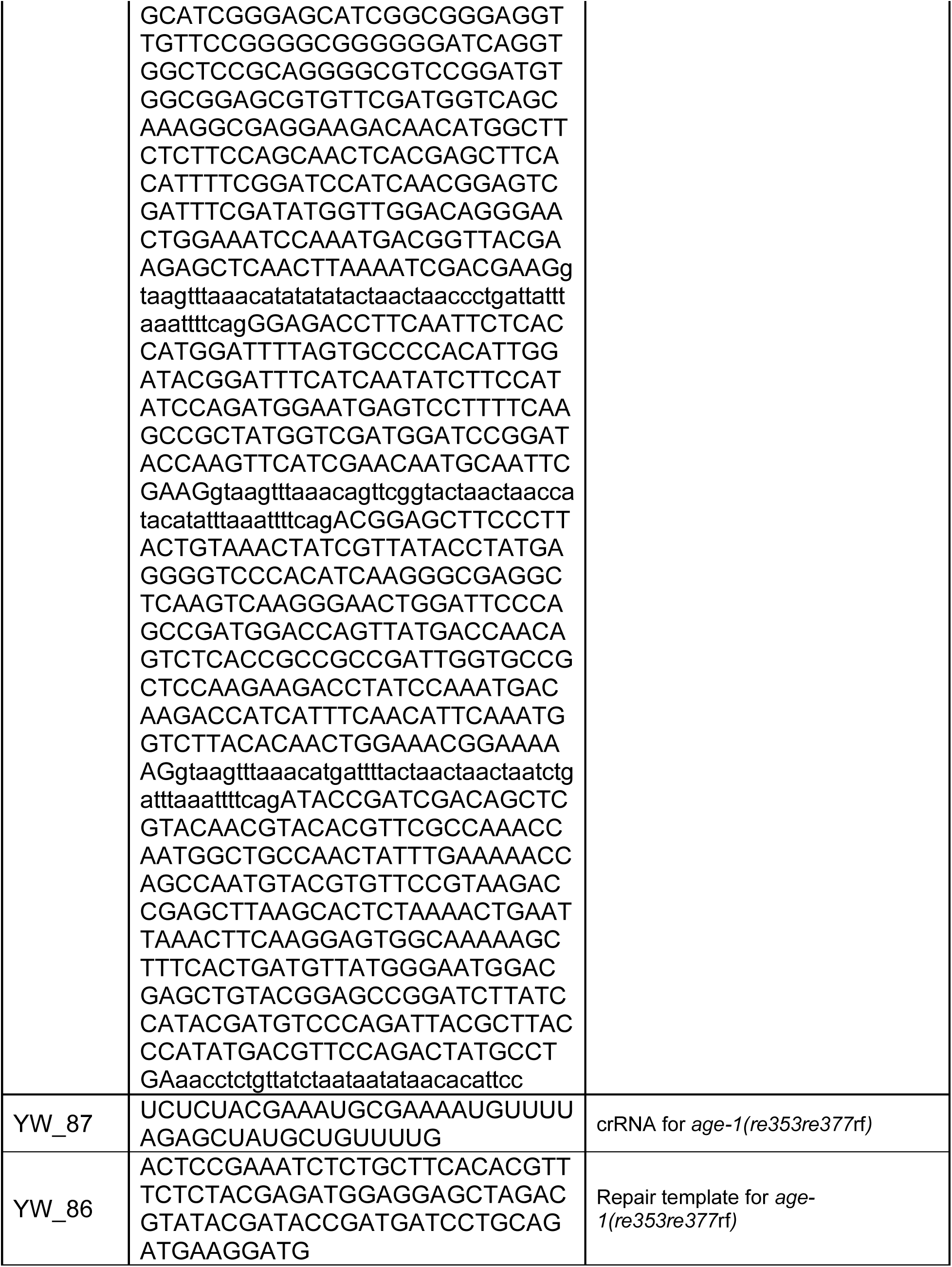
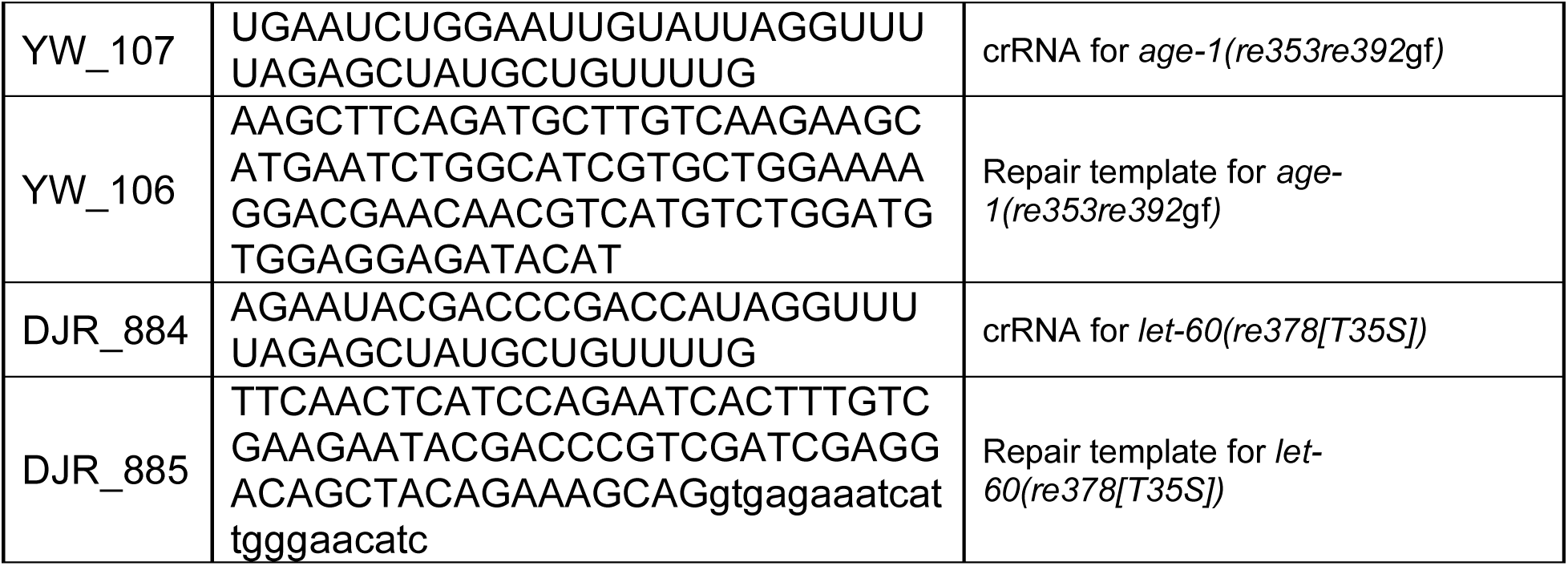
CRISPR reagents.

